# A deep single cell mass cytometry approach to capture canonical and noncanonical cell cycle states

**DOI:** 10.1101/2025.07.08.663243

**Authors:** Meelad Amouzgar, Patricia Favaro, Daniel Ho, Trevor Bruce, Sean C. Bendall

## Abstract

The cell cycle (CC) is involved in diverse cell processes like cell differentiation, immune cell expansion, and tumorigenesis but current single-cell (sc) strategies study CC as: (i) coarse phases, (ii) are based on transcriptomic data, (iii) leverage imaging modalities that are not extensible outside adherent cell culture, or (iv) do not enable high-throughput, single-cell multiplexing. To solve this, we developed an expanded, metal-tagged antibody approach for Mass Cytometry (MC) with 48 CC-related molecules that more deeply phenotypes the diversity of scCC states. Using cytometry by time of flight (CyTOF), we quantified CC states across various suspension and adherent cell lines and stimulated primary human T cells. Our approach captures the diversity of scCC states, including atypical molecular signatures that do not meet the canonical definitions of CC phases in the literature. Multiplexing with pharmacologically-induced CC arrest reveals that drug-induced CC perturbations can exacerbate reported noncanonical states and induce previously unobserved states. Notably, cells escaping CC inhibitor action in primary cells demonstrated more aberrant CC states compared to their untreated counterparts. Overall, our approach enables deeper phenotyping of CC biology that generalizes to diverse cell systems and can simultaneously multiplex experimental perturbation & disease systems, or integrate with other MC measurement platforms.

## Introduction

Regulated cell division via cell cycle (CC) is a pervasive and ubiquitous biological process in both health and disease (1). Here, the cellular decision to proliferate is under tight molecular control to maintain organismal homeostasis and rapidly expand immune cells while preventing unregulated expansion (2). A central challenge of quantifying and assigning CC state is that cycling cells can be rare and heterogeneous - thus single cell approaches are often employed to quantify CC states. The detection of proliferative signatures is further convoluted by differences in cell types, lineages, disease, and drug treatment, where intrinsic differences to CC architecture or extrinsic disruptions to CC molecular state may not be captured by low-dimensional measurement strategies (3).

Fluorescence flow cytometry (FCM) can discriminate the four major cell cycle (CC) phases of G0, G1, S, G2, and M using measurements of DNA content (i.e. PI, 7AAD), nucleotide incorporation (i.e., BrdU, EdU), protein expression and phosphorylation (i.e., cyclin B, phospho-H3) (4,5). The advent of next-generation single-cell (sc) approaches, such as Cytometry by Time of Flight (CyTOF) Mass Cytometry (MC) have enabled quantitative multiplexing beyond 45 molecular targets per cell facilitating both regulatory molecular measurements and deep phenotyping of cell identity (6,7). Our prior work using MC includes a combination of Cyclin B1, phosphorylated-Histone H3 (pHH3, Ser28), and incorporation of 5-iodo-2′-deoxyuridine (IdU) allowing the manual annotation of G0/G1, S, G2, and M phases (8). More granular separation of CC phases has been further achieved with the addition of phosphorylated retinoblastoma (pRb), Geminin, CDT1, and PLK1 to discriminate between G0 and G1, and split G1 and G2 phases into early and late stages (9,10).

Despite these advances in cell cycle phase discretization, analysis methods are still overly-manual, relying on gating strategies to categorize and quantify cell bins. This approach fails to recognize the continuous nature of cell cycle progression, or how cells can occupy unique normal and dysfunctional CC states depending on intrinsic cell processes as well as extrinsic perturbations like drug inhibitors that induce CC arrest via mechanisms such as CDK inhibition, the spindle checkpoint, or p53 activation (11). While such advancements in quantifying CC aberrancy have been approached with live-cell microscopy to describe the molecular architecture during hypomitogenic and replication stress, an advantage of next-generation single cell (sc) approaches, like MC, is they are well-suited for economical, high-throughput analysis of dissociated cell systems in combination with barcoding strategies for multiplexing experimental samples, enabling deeper (i.e. higher throughput and multiplexing) characterization of CC states beyond manual gating of CC phases (12–15,15). The global dynamics of molecules regulating CC are often conserved across different cell systems but the exact abundance and patterning of molecules required for cell cycle progression can differ depending on factors like cell size, genome size, replication speed, cell line origins, and cell extrinsic molecular factors (16–18). Furthermore, disease and perturbation can disrupt the molecular patterns that define canonical CC, inducing noncanonical CC states like CyclinD1 loss in G2 phase, relicensing of DNA replication by CDT1 overexpression during G2 phase, and tetraploidy (12,19–21). High-throughput low-dimensional strategies may fail to capture or deeply characterize both canonical and noncanonical CC states.

The high throughput, parallel quantification capabilities of scMC has allowed phenotyping of molecular states during complex biological processes in primary cells, such as TCR stimulation and T cell expansion, as well as chromatin modifications across immune cells (22,23). To better understand the diversity of scCC states using MC, we further multiplex existing MC panels to expand CC-related measurements. Using this, we characterize the diversity of CC states across five cell lines commonly used in the literature as well as stimulated and expanded primary human T cells. We further study the consequences of CC perturbation on scCC states using DNA synthesis inhibitors, microtubule destabilizing agents, and CDK4/6 inhibition in Jurkat cells as well as a breadth of CDK inhibitors in primary human T cells to quantify the patterns of scCC aberrancies in arrested and escaped cells. Using our scCC platform in primary human T cells, we reveal the CC state similarities in drug perturbation between different CDK inhibitors, the consequences of CDK inhibition on the diversity of scCC states, and expose that cells escaping CC arrest occupy aberrant CC states not observed canonically. This new single cell assay serves as an additional MC tool that can be integrated with other MC platforms like scMEP for metabolism, EpiTOF for chromatin marks, cell phenotyping, cell differentiation, hematologic cancer cell, and other measurements to tease the pervasive crosstalk between CC and other systems (22–26).

## Results

### A single cell proteomic module for deep molecular typing of cell cycle biology

The cell cycle is a highly conserved biological process with tightly regulated checkpoints finely orchestrated by CC protein abundance and post-translational modifications (PTMs). This molecular machinery ensures accurate DNA replication and chromosome segregation during cell division and prevents premature S- and M-phase entry from the gap phases (27). To date, CC quantification by single cell analysis has focused on quantification of cells residing in individual phases using landmark reporters (9). However, recent studies demonstrate that cells can occupy diverse CC states captured with multivariate molecular signatures (12). For example, gold standard markers of proliferation Ki67 and phosphorylated Rb are dynamically expressed and can be lowly abundant in cells with other cycling signatures, like active DNA replication (S phase) defined by IdU labeling or high cyclin B abundance (G2 phase) (**Figure 1A**) (28). For this reason, low-dimensional landmark reporters fail to capture more granular CC states in exact phases, and may not accurately quantify the molecular effects of CC perturbation. To establish a solution for this we created an expanded, metal-tagged antibody approach that quantifies the abundance of an increasingly diverse number of CC-related molecules including: cyclins that promote CC progression, CC checkpoint inhibitors, DNA licensing factors, PTMs of Rb and histone proteins that control proliferation and mitotic entry, nucleotide incorporation during S-phase, DNA content, chromatin state, and CC regulators (<u>Table S1</u>). Our scCC approach includes three sets of molecular features: (i) ‘minimal’ includes protein and phospho-protein targets that directly control cell cycle checkpoints and progression, (ii) ‘core’ includes the minimal CC molecular targets paired with measurements of DNA content and replication such as DNA intercalators and IdU incorporation, and (iii) ‘complete’ includes a wide array of measured CC-related molecules including transcription factors, chromatin state, and other CC regulators (**Figure 1B**). These targets were categorized into each panel based on both variance associated with separating cell cycle phases and functional characteristics reported in the literature (**Figure 1B, S1A-C**). This panel can be further customized to include new targets. We additionally recommend a combined experimental and *in silico* cleaning strategy for debris, doublet, and pre-apoptotic cell elimination that generalizes to any experimental system using CyTOF (**Figure 1C-D**). With our scCC platform, we further characterize CC heterogeneity in diverse, suspension and adherent cell lines (**Figure 1E-G)**. We establish a core CC molecular panel with breadth to directly measure diverse CC states for combinatorial sc systems analysis with other MC panels (**Figure 1E**). We then extend our platform to multiplexed perturbation experiments and quantify the sc consequences of a variety of CC inhibitors in both cell lines and primary human T cells (**Figure 1F**). Using a graph connectivity approach to quantify cell diversity, our platform for deeper CC profiling captures increasingly more diverse CC states as a function of additional CC-specific features across all cell lines (**Figure 1G-I, see methods**). Notably, there were cell-line dependent differences in CC diversity captured by the panel in both the average diversity as well as the increase in diversity with additional features.

**Figure 1.**
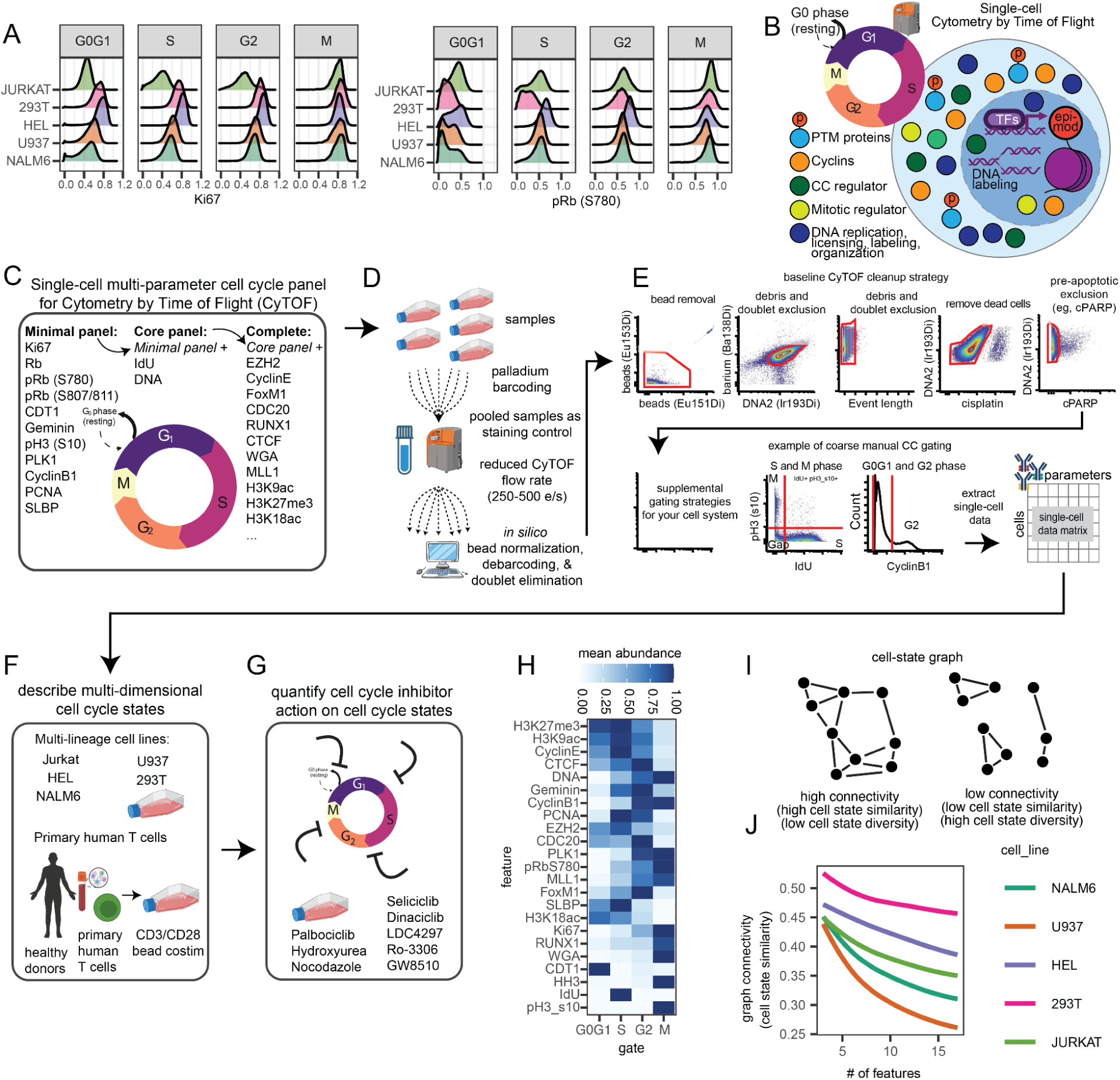
A revised targeted approach to capture granular cell cycle information. **(A)** Example markers with CC distribution differences across cell lines facetted by coarse manually-defined groups. **(B)** Cartoon of different types of CC-related molecules. **(C)** Cartoon schematic for scMC panel of CC-related molecular targets for the minimal, core, and complete panels (see <u>Table S1</u> for all targets). **(D)** Cartoon schematic of controlled experimental multiplexing strategy for doublet reduction using palladium barcoding. **(E)** Recommended minimal gating strategy for data cleanup and example of coarse manual CC gating strategy. **(F)** scCC panel applied to cell lines with different celltype identities and primary human T cells in proliferating and **(G)** CC perturbation settings. **(H)** Heatmap of CC-relevant markers used in cell line analysis (n=3). **(I)** Schematic of graph-based approach to quantify cell state similarity using edge density. Adjacency matrix computed using a cosine distance with a threshold of 0.5. **(J)** Mean connectivity (cell state diversity) as a function of # of features along all possible combinations of features in each cell line.

### Dimensionality Reduction Reliably Captures Cell Cycle Phase

To understand the CC molecular expression across different cell systems, we applied our molecular panel to a variety of cell line types that includes both adherent and suspension cells: NALM6, U937, HEL, 293T, and JURKAT cell lines, as well as primary human T cells. Given that different cell systems can have variable CC dynamics, we first sought to evaluate the diversity of CC states in these cell lines (**Figure 2A-B**) (29). We leverage multiplexed palladium barcoding to eliminate procedural variation between samples & reduce doublets, simultaneously process and stain cells for fair molecular comparisons across cell lines, and perform live singlet acquisition using event length, barium, DNA, and standard CC gating practices to remove doublets, debris, dead, and pre-apoptotic cells (10,24,30).

**Figure 2.**
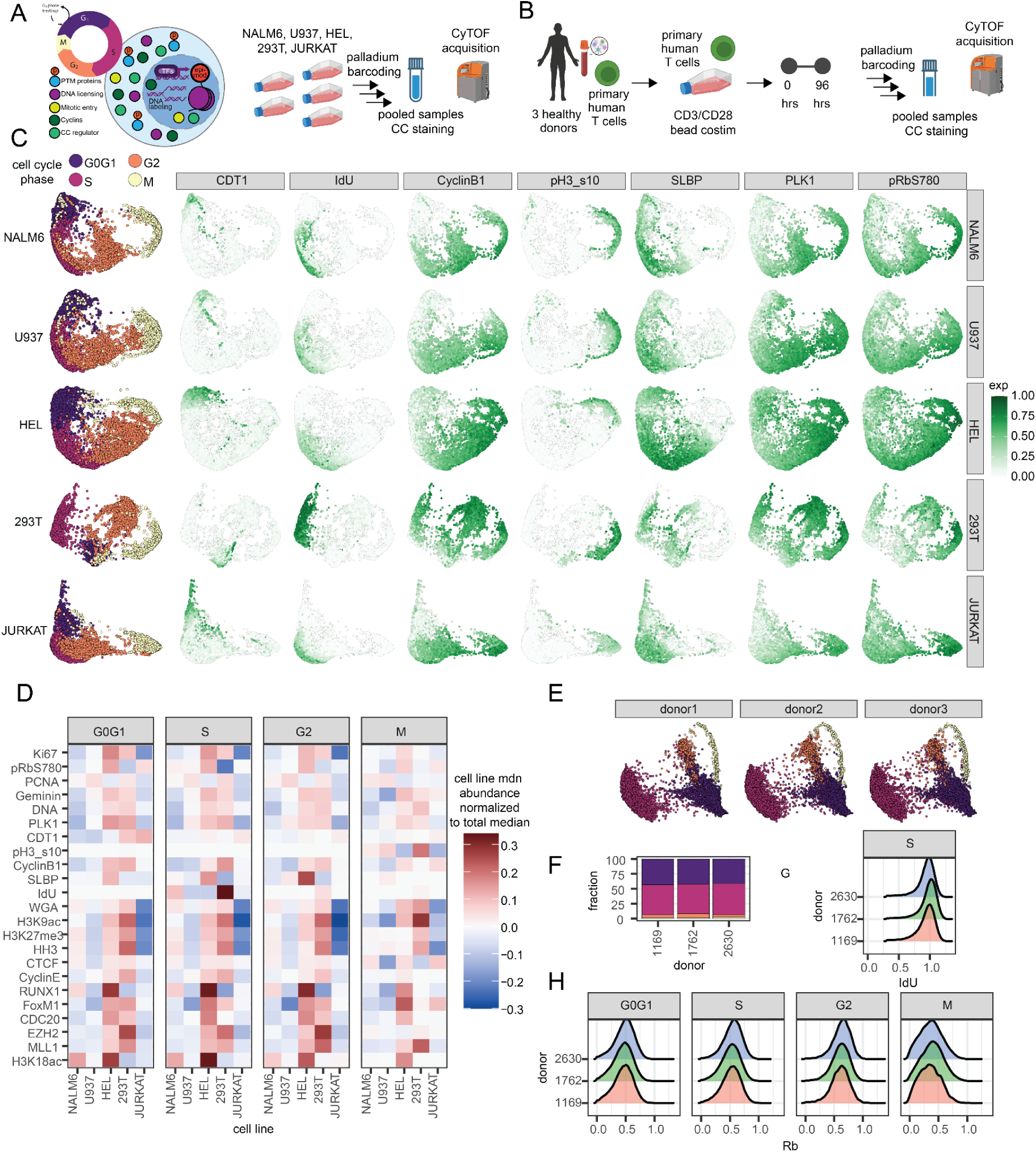
CC embeddings and variance of CC information. **(A)** Experimental design for CC analysis by CyTOF across palladium-barcoded cell lines (n=3 each) pooled from multiple cultures. **(B)** Experimental design for ex vivo primary human T cell activation using CD3/CD28 dynabead stimulation across 3 unique donors (**C**) Dimensionality reduction of cell lines and example markers from CC panel using PHATE. (**D**) Median molecular abundance of CC targets for each cell line normalized to median across all cell lines, each equally sampled. General CC phases are defined using CyclinB1, IdU, pH3(s10). (**E**) CC embedding molecular panel to CD3/CD28 dynabead stimulated primary human T cells from 3 donors. (**F**) Cell cycle phase fractions after 4 days of stimulation. (**G**) Donor variability in DNA replication measured by IdU incorporation during S phase and (**H**) Rb expression across manually-grouped phases.

Independent dimensionality reduction of each cell line with PHATE captures expected separation of the major G0, G1, S, G2, and M CC phases defined by a classical gating strategy using IdU incorporation for S-phase, pH3 (Ser10), and CyclinB1 (**Figure 2C**) (31). We observe consistent patterns of CC molecules that capture CC progression in each cell line, such as higher PLK1 and pRb (S780) expression in S, G2, and M phases, a reduction of SLBP as cells exit from S through G2, and CDT1 licensing for DNA replication primarily restricted to G0G1 cells (**Figure 2C**). Notably, the individual cell line CC embeddings were organized differently from each other - with NALM6 and U937 consistently having more similar embedding structures, and 293T cells having the most different - along with the patterning and abundance of CC molecules suggesting differences in CC molecules across cell lines.

### Variability in Cell Cycle Molecules Across Phases Is Cell-type Specific

Considering these differences in CC embeddings and the shifted molecular distributions across cell lines, we compared the mean expression of each cell line against the total mean of all cells for each molecular target from a random, balanced sampling of each cell line to quantify cell line variance (**Figure 2D**). We observed a trend of HEL and 293T cells having a globally higher abundance of CC-relevant molecules such as Ki67, PLK1, CyclinB1, and Geminin compared to NALM6, U937, and JURKAT cells. Notably, we observe the same patterns in WGA abundance and total histone content - proxies of cell size (24,32,33). HEL and 293T cells tend to be larger (∼12-15 um) while the leukemic cell lines Jurkat and NALM6 cells are smaller (11.5 um) (34,35). However, U937 - a pro-monocytic cell line - is larger (∼13.29 um) but does not express as much global CC molecule abundance as HEL and 293T (36). Taken together, this suggests that while cell size may explain some of these differences unveiled by our scCC approach, the cell type origins and underlying origins of these cell lines likely explain the majority of variability in CC molecules across these systems.

With variation between cell lines evident, we were curious whether similar primary human cells from different individuals could exhibit similar traits. To investigate this, we queried primary human T cells from 3 different healthy individuals with our CC panel. Using primary human T cells sampled 96 hours after *ex vivo* CD3/CD28 bead stimulation from 3 donors, we observed topologically similar CC embeddings between individuals, and similar to those seen for the cell lines (**Figure 2E, Figure S3A**). CC phase occupancy defined by manual grouping averaged about 42% G0G1, 51% S, 5.1% G2 and 1.8% M across 3 donors, with little differences between donors (**Figure 2F**). Considering there is known human variation in molecules across different cell systems, we did not observe differences in donor-dependent variation in CC-related molecule abundance like IdU, CyclinB1, pH3 (s10), and Rb. (**Figure 2G-H, Figure S3A).** Our approach also captures expected abundance patterns, such as SLBP abundance higher in S, and a small subset of SLBP high cells in G2 as it undergoes rapid degradation upon G2 entry. As well as PLK1, Ki67, and PCNA abundance increasing with CC progression, and pRb (S780) expression increasing in both G0G1 and S phase (**Figure S3A**). Taken together, our CC panel generalizes to diverse cell lines as well as primary human cells, with measurable differences in CC molecular abundance between cell lines despite the CC being a conserved molecular program intrinsically tied to a cell’s biology.

### Multivariate CC quantification reveals parallel but distinct CC patterns between cell lines

Considering the variance of CC molecule abundance patterns we observed across cell lines was robust across replicates (**Figure S2A, S4A-B**), we sought to leverage our deep CC phenotyping panel to further integrate and understand the molecular abundances in aggregate. Pairwise comparison of cell lines across all replicates identified NALM6 and JURKAT cells with the fewer CC-related molecular differences **(Figure S4C)**. CC embeddings showed a coordinated separation of CC phases with clear cell line-dependent patterns (**Figure 3A-B**). Single-cell data integration methods like Harmony are often used to correct batch or biological confounders, yet these approaches come at a tradeoff between batch removal and bio-conservation (37,38). Employing Harmony to correct cell line differences adequately integrated cell line differences, however, CC phase purity analysis using FlowSOM for multivariate unsupervised clustering showed that while S phase cells observed an improvement in cluster purity after correction, G0G1 and G2 purity decreased (**Figure S4C-D**). Considering the tradeoff with using correction algorithms leading to the removal of biological signal or insertion of non-biological signal, and the lack of systematic single-cell comparison of CC states between diverse cell lines, we bypassed biological batch correction and further studied these cell line differences (39,40).

**Figure 3.**
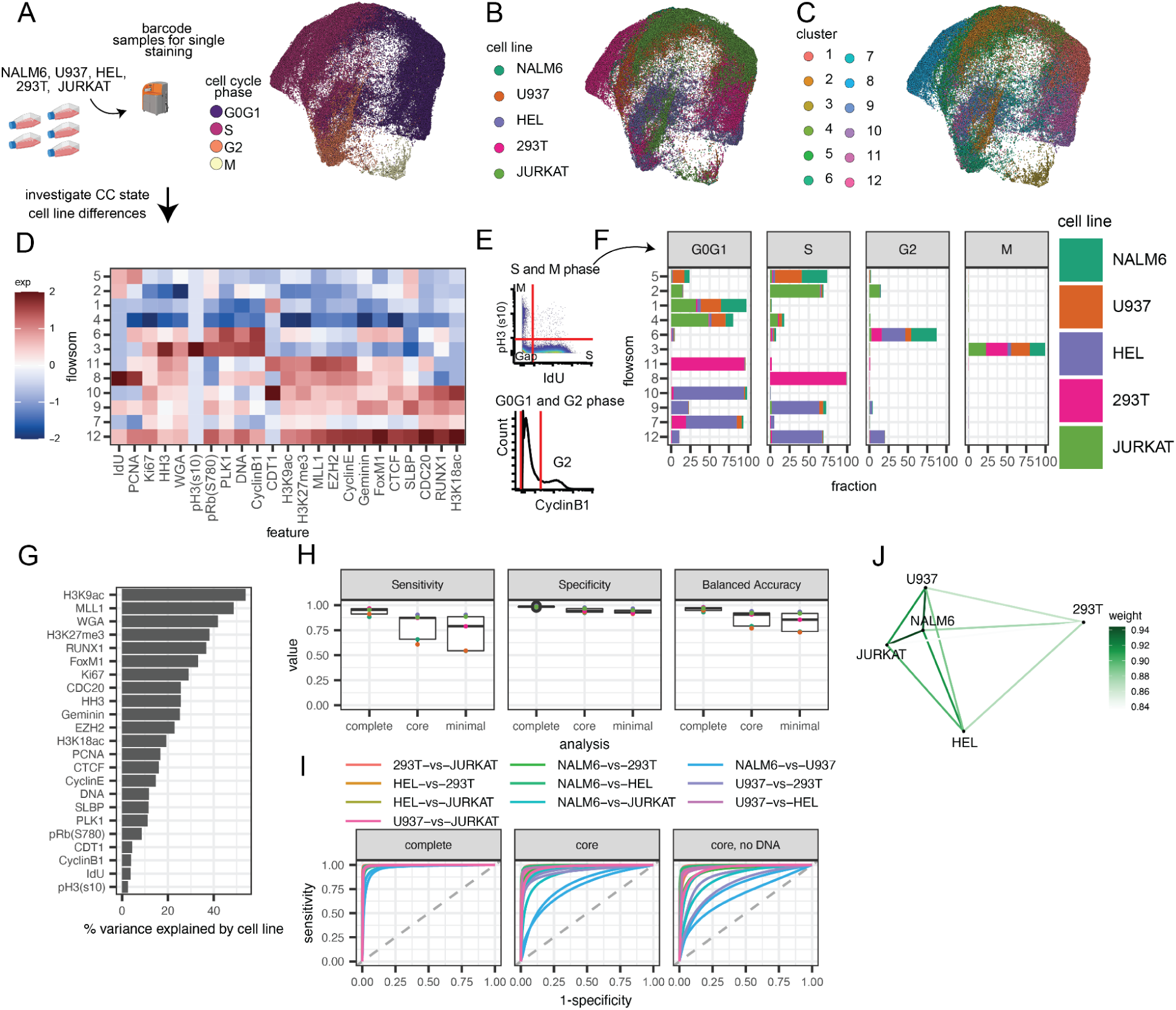
Integrated multivariate CC analysis reveals differential cell-line CC states. **(A)** CC embedding of molecular targets equally sampled for each cell line pooled from multiple cultures, and z-scored across all cells in aggregate (n=4 each across two experiments, showing 1 replicate each). Colored by **(A)** cell cycle group, **(B)** cell line, **(C)** and flowsom cluster. **(D)** Heatmap of CC molecular target abundance across clusters. **(E)** Current gold-standard gating strategy for identifying G0G1, G2, DNA replicating (S) and Mitotic (M) cells. **(F)** Fraction of cell lines in each cluster faceted by CC phase. **(G)** ANOVA results measuring % variance explained by cell line identity for CC molecular targets. **(H)** Sensitivity, Specificity, and Accuracy results for classifying cell line identity in a test set using multinomial logit regression with the complete, core, and minimal CC panels. **(I)** ROC curve using each panel. **(J)** Analysis of different CC molecular correlation patterns between each cell line showing similarity between cell lines using the core panel.

Deeper probing of unharmonized clustering shows that clusters 2,8,9,10,11,and 12 capture cell lines with CC phase specificity, except for clusters 9 and 12 which have mixtures of G0G1/S or G0G1/S/G2 phases. Clusters (1,3,4,5,6,7) capture mixtures of cell lines with G0G1, S, and G2 phases defined by IdU, pH3(s10), and CyclinB1 expression. Interestingly, cluster 3 captured the majority of mitotic cells regardless of cell line status **(Figure 3C-F)**, which is consistent with previous studies looking at multivariate molecular measurements of chromatin state across CC phases in different cell lines (41). Clusters (5,6,8,9,12) with more proliferative phases (S/G2/M) had generally greater molecular abundance of key molecules like Ki67, PCNA, and Geminin but clusters 2 is an interesting exception capturing cells with active DNA replication (IdU+) cells that are Ki67 low. CDT1 positivity in clusters 1 and 10 indicates replication licensing before S phase but cluster 10 primarily consists of HEL cells with increased abundance of transcription factors like FoxM1 and CTCF, as well as SLBP, while cluster 1 is has lower abundance of most CC-related molecules and includes diverse cell lines. High SLBP abundance in cluster 4 indicates deep progression into S phase as SLBP coordinates histone synthesis with DNA replication but again, this cluster is primarily detected in HEL and 293T cells. The majority of G2 cells are included in cluster 6, which has expected high expression of CyclinB1, PLK1, pRb (s780), and Ki67 but small fractions of G2 cells also appear in other clusters that are low in these molecules typically expressed in G2 cells, such as cluster 2. SLBP loss is observed in clusters with larger G2 fractions (6,12), which is expected since SLBP degrades rapidly during the S/G2 transition (42). In summary, cell line variance captured by the scMC CC platform is a convoluting factor in understanding cell cycle states.

Considering cell line variance confounds CC states with overclustering, we simplified the cluster task by reducing to four clusters. There were mostly pure clusters of G2 and mitotic cells in clusters 3 and 4 while G1, S, and G2 phases were mixed between clusters 1 and 2 **(Figure S4E)**. These results suggest that whether under-clustering or over-clustering, basal cell line differences in CC states complicate more granular CC analysis, and that cell line variance is a strong contributor to CC state diversity. CC state integration may be necessary for experimental designs seeking to combine different systems like cell lines. But the loss in signal associated with integration, the observed purity differences between CC phases, and the mixing of CC phase and cell line identities all demonstrate that our scCC approach captures disparate CC molecular programs that are unique to each cell line.

### Machine learning models quantify differences in CC molecular abundance across cell lines

Considering the strong multivariate differences in CC molecular abundance across the cell lines, we sought to further quantify these molecular differences using interpretable univariate and multivariate statistical models that are balanced for each cell line in the training data. To estimate the contribution of each molecular target in separating the cell lines, we performed Analysis of Variance (ANOVA) of simple linear regression models to predict molecular abundance from cell line identity **(Figure 3G)**. Molecules capturing chromatin state (H3K9ac, MLL1, H3K27me3) and cell size (WGA) as well as molecular regulators that are not necessarily CC-specific (RUNX1) had stronger associations explaining cell line differences (**Figure 3H**) (24,32). However, classical CC molecules like Ki67, CDC20, and PCNA also explained considerable variation. Consistent with the clustering results and literature analysis of chromatin state, pH3 (s10) poorly explained variance across cell lines, again suggesting that the phospho-chromatin state of mitotic cells are similar to each other (41).

To further quantify the differences in CC states in the cell lines, we trained a multinomial logistic regression model and evaluated their performance predicting cell lines using the three CC feature sets from Figure 1: (i) ‘minimal’, (ii) ‘core’, and (iii) ‘complete’ **(molecules labeled in Table S1)**. Multivariate models predicting cell line outcome using all CC-relevant targets performs well with over 82% sensitivity, 97% specificity and 96% accuracy across all cell lines indicating robust cell line predictions (**Figure 3I-J**). However, the removal of chromatin state, cell size, and molecular regulators in the core panel reduces predictive performance, and the additional removal of DNA-related measurements in the minimal panel further reduces performance. Predictions using core cell cycle markers achieve 75% accuracy, indicating that there remains fundamental differences in CC-specific markers across cell lines **(Figure 3I)**. Pairwise comparison of cell line predictions in the core and minimal panels further shows that comparisons such as NALM6-vs-U937, U937-vs-293T, NALM6-JURKAT have poorer model performance, suggesting greater similarity between CC states due to less differences in CC molecular abundance between these cell lines **(Figure 3J)**. Correlation network analysis (CNA) similarly suggests disparities in multivariate abundance patterns between cell lines, where JURKAT, NALM6, and U937 are most similar to each other. Notably, 293T - an adherent cell line - has the most different CC co-abundance pattern compared to the other suspension cell lines.(**Figure 3I-J**). Hierarchical clustering of CC abundance across replicates supports these results **(Figure S4B)**.

We further investigated whether these CC differences are explained by single nucleotide variant (SNV) or copy number variant (CNV) profiles of cell cycle-related genes using DepMap(43) published information for Jurkat, HEL, U-937, and NALM-6 (293T data was unavailable). Variant profiles were in agreement with the cell line heterogeneity observed in our scCC MC platform, though the majority of gene variants do not intersect with molecules probed in our platform (except PCNA) **(Figure S4G-I, Table S2-3)**. These results suggest that genetic sensitivity to cell cycle-related variants may have gene-level biases based on cell line origins or evolutionary origins, which are being detected indirectly by our scCC panel. While the CC is a generally conserved evolutionary program across different cell systems, we demonstrate that our approach can probe deeper into the diversity of cell states and reveal the subtle differences across similar but disparate CC molecular programs across these diverse cell lines.

### Single cell analysis across cell lines decouples canonical and noncanonical CC states

We demonstrated that deeper CC probing using our approach revealed the diversity of CC states across cell lines. CC states are often described by canonical rules like Ki67 expression as a marker for proliferative cells in S, G2, or M phase, and CDT1 licensing for DNA replication during G1. However, CC aberrancies are a hallmark of diseases like cancer, and these noncanonical CC states are reported across diverse, transformed cell lines (44). Examples of noncanonical CC states induced by CC perturbation can include CDT1 overexpression in G2 phase leading to relicensing of DNA replication if unregulated by Geminin, failure to degrade SLBP during G2 leading to genotoxic stress, chromosome instability from various consequences such as loss of PLK1 during mitosis leading to mitotic slippage and cellular senescence, Ki67 loss or dephosphorylation of Rb while cycling, and other mechanisms that disrupt CC progression. Noncanonical CC states have been reported in cells experiencing DNA damage, mitotic infidelity, or disruption of CC regulators (19–21,28,45,46). We observe that our scMC approach captures these noncanonical CC states without perturbation, such as low Ki67 abundance in S phase cells actively replicating DNA, CDT1 expression during G2, high pRb (S780) or PLK1 expression during G0G1, low pRb (S780) in G2, and additional noncanonical cell states (**Figure S2**) (47). To label noncanonical CC states, we discretized cells into canonical and noncanonical groups based on rules for canonical CC states found in the literature for features in our core panel and observed variable fractions of noncanonical CC states between cell lines and features (**Table S4**). For example, canonical CC states with clear proliferative signatures are expected to express Ki67, or DNA licensing (CDT1) should not be abundant after S-phase, and thus deviancy from these cell states could indicate CC aberrancy. All cell lines had a subset of cells actively replicating DNA (IdU+) while Ki67 low, CDT1-expressing cells in G2, and G2 cells that were pRb (S780) low, particularly in 293T cells (**Figure S5A-C**). Manually discretized noncanonical CC phenotypes were on average 23.9% of cells (NALM6, 14.6%; U937, 16.6%; HEL, 26.5%; 293T, 36.7%, and JURKAT, 25.3%) (**Figure 4A**). Mahalanobis distance–a multivariate measure of each point (cell) from its population centroid, where larger magnitudes indicate further distance–of all cells in each phase was higher for noncanonical cells compared to canonical cells in the same phase, suggesting these grouped cells are distinct from the average cell in each phase, with greater median aberrancy for noncanonical cells in S and M phase compared to their matched canonical counterparts (**Figure 4B**). Considering that noncanonical CC states can be representative of perturbed CC progression or altered CC states, we asked whether including only canonical or noncanonical cells as input into dimensionality reduction would more accurately capture CC progression. Indeed, we observe improved topologically circular CC embeddings when excluding noncanonical cells, and aberrant topologies when excluding canonical cells (**Figure 4C**). By more deeply measuring CC-related molecules at the single-cell level, we can reconstruct CC as historically learned through bulk assays as well as parse the diversity of CC states that may be over-generalized if studied through the lens of coarse CC phase annotation.

**Figure 4.**
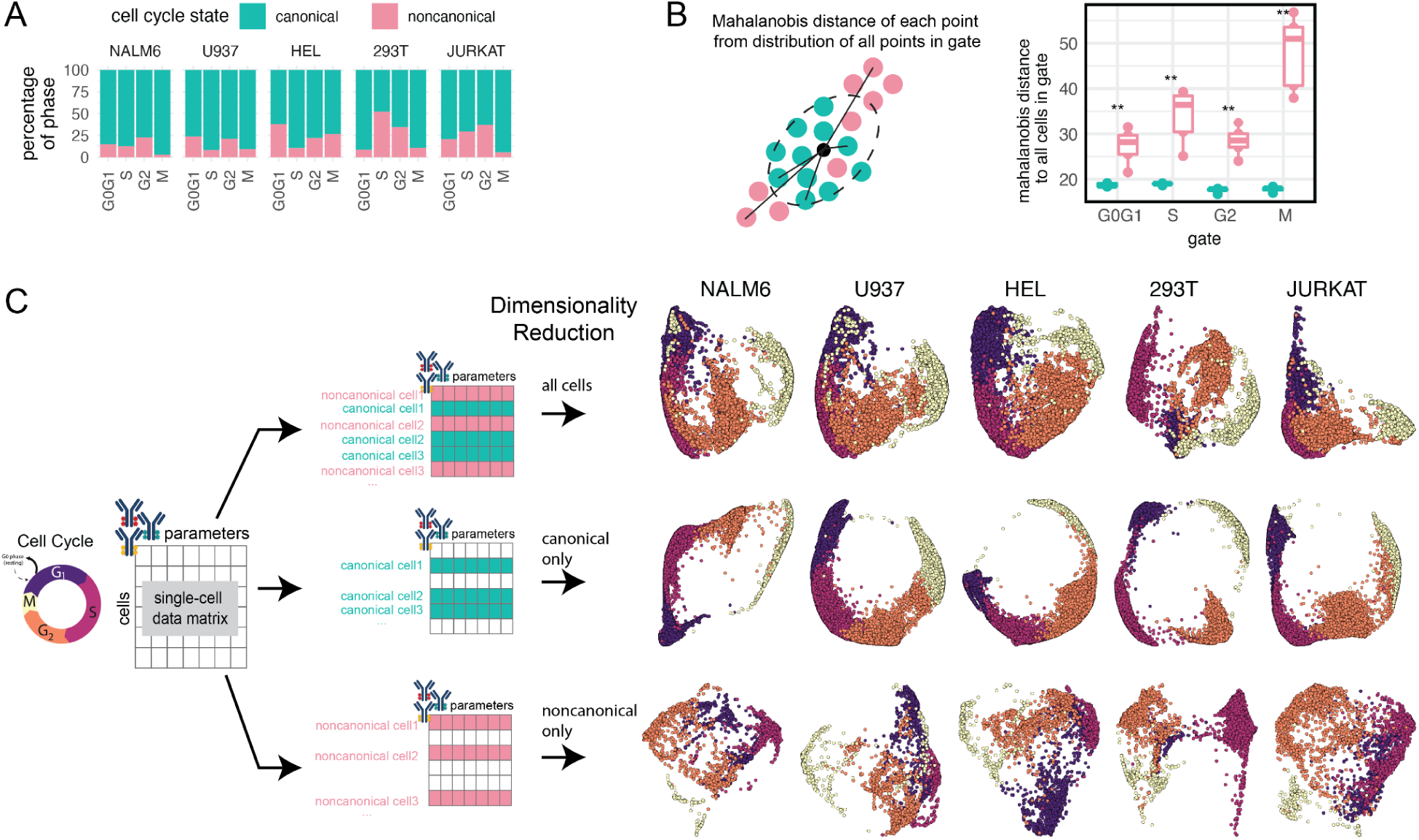
Multi-parameter CC analysis captures canonical and noncanonical single-cell CC states in diverse cell lines. **(A)** Estimated fraction of manually discretized noncanonical cells in each cell line and CC phase based on a manual discretization strategy. **(B)** Mahalanobis distance analysis comparing noncanonical cells and canonical cells to all cells (wilcoxon rank sum test, -p values: * <=0.05, **<=0.01). **(C)** CC embedding constructed on cells with both canonical and noncanonical, canonical-only, and noncanonical-only CC states using core panel features.

### Cell cycle perturbation induces new noncanonical cell cycle states

While we quantified fractions of noncanonical CC states that reflect previously reported CC aberrancies, prior studies have also shown that pharmacologic inhibition disturbs the molecular state of CC progression (48–50). Similarly, we used CC inhibitor assays to pharmacologically target different CC stages and study whether we can: (1) induce and capture known and new noncanonical scCC states; and (2) quantify inhibitor action. We treated Jurkat T-cells with three different CC synchronization drugs for ∼18 hours and evaluated protein biosynthesis before harvesting cells for CyTOF using our platform after multiplexing samples with barcoding and removing dead and preapoptic cells *in silico* (**Figure 5A, S6A, Table S5**). We used Palbociclib (PALBO), a CDK4/6 inhibitor that primarily arrests cells in G0G1; Hydroxyurea (HU), an inhibitor of ribonucleotide reductase (RNR) that reduces the dNTP pool to slow down DNA polymerase movement at replication forks and activate the S-phase checkpoint; and Nocodazole (NOC), a microtubule disrupting agent that binds to beta-tubulin and arrests cells at the G2M checkpoint, which can also induce mitotic slippage into G1 (48–50)

**Figure 5.**
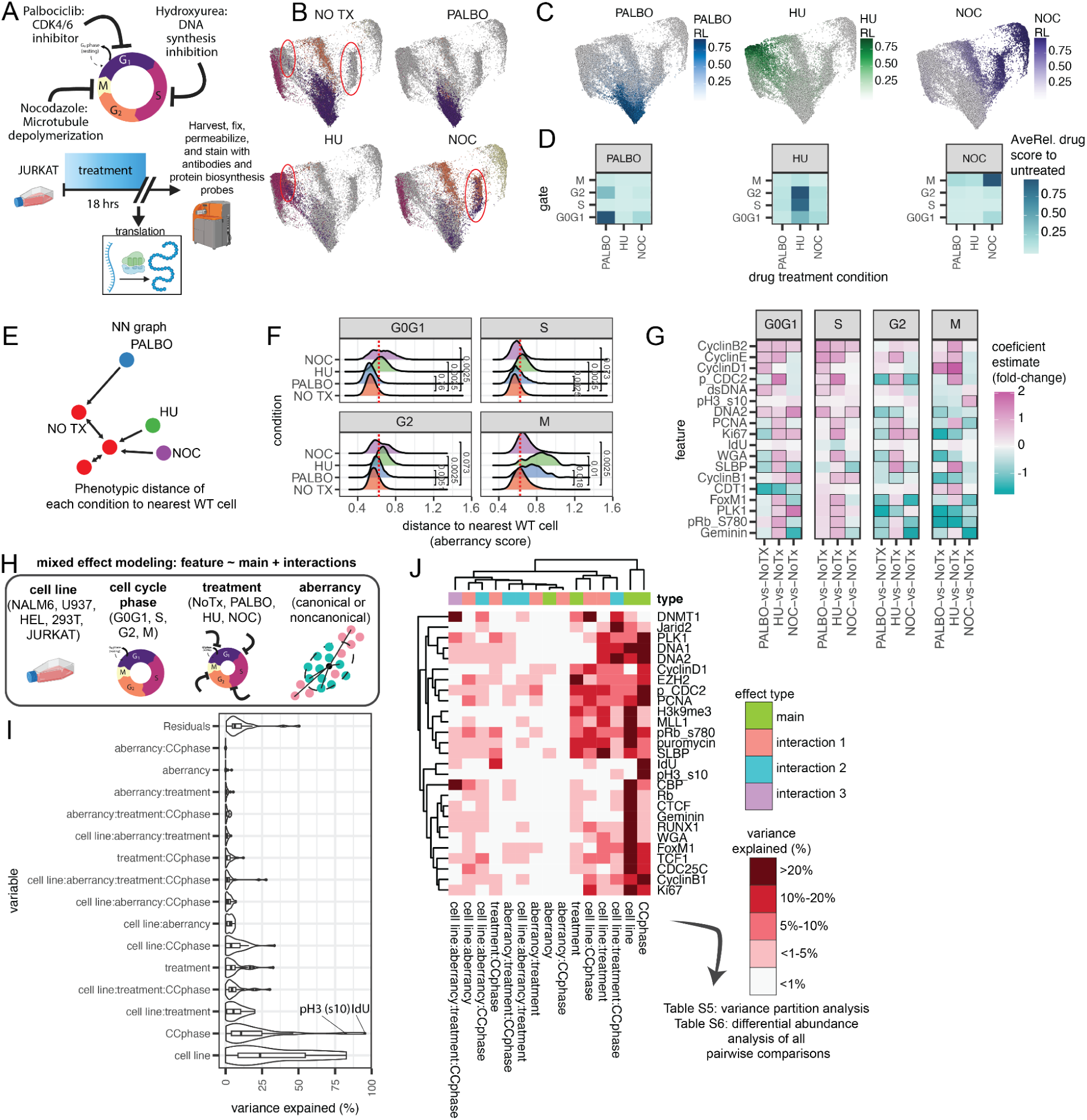
CC inhibitors induce non-canonical states. **(A)** Cartoon schematic of pharmacologic inhibition in JURKAT cells (n=5 from 3 experiments). **(B)** CC embeddings for each drug with all cells in gray background, highlighting induced CC states (1 representative sample). **(C)** CC embeddings with perturbation score quantified using MELD. **(D)** Median perturbation score for each group in each drug condition. **(E)** NN analysis cartoon schematic quantifying phenotypic CC distance from untreated cells. **(F)** Distribution of CC distance, dotted line indicates a manually-defined threshold for canonical cell identity based on the WT distribution across. Wilcoxon mean test on mean aberrancy from each sample. **(G)** Heatmap results comparing drug treatment effects to NoTx (black box indicates significance defined by padj<=0.1 and absolute coefficient estimate >= 0.5). **(H)** Cartoon schematic for single-cell variance partitioning analysis across cell lines, cell cycle phase, drug treatment, and aberrancy (n=36 unique samples, 3,544,481 single-cell measurements: 3 replicates each cell line, 3 replicates of each drug treatment in Jurkat cells, 1 replicate of each drug treatment in NALM6, U937, HEL, 293T cells). **(I)** Results of statistical modeling of differences of noncanonical variables defined by NN phenotypic distance to untreated cells. **(J)** Binned percentage of variance explained by each main or interaction effect for each molecular target.

Drug-treated cells piled-up at expected CC phases, and CC embeddings suggested drug-treated cells occupy phenotypically distinct states from untreated (NO TX) cells (**Figure 5B**, red circles). We further evaluated the inhibitory effects of PALBO, HU, and NOC at single-cell resolution by quantifying the perturbative action on CC relative to the untreated condition using MELD–a computational algorithm that uses graph signaling processing techniques to measure the sample-associated relative likelihood between experimental groups (eg, treatments) (**Figure 5D, S6B**) (51). PALBO inhibitor action is primarily observed in G0G1 and a small subset of G2-like cells. HU inhibitor action is observed in G2, S, and G0G1 cells. NOC inhibitor action is observed in M and G0G1 cells (**Figure 5D**). Inhibitor action on CC states is largely unique to each inhibitor, though some cells with G0G1 CC states induced by PALBO are also observed in NOC-treated cells, and vice-versa. Scoring CC phase enrichment defined by manual groups further quantified inhibition action in treatment conditions, and our platform clearly captured noncanonical CC states induced by drug treatment in arrested cells as well as cells that escaped arrest (**Figure S6C**).

We previously quantified noncanonical CC states using manual discretization but this relies on coarse gating practices. To more deeply characterize induced noncanonical CC states relative to an unperturbed system, we used nearest neighbor (NN) analysis to quantify the multivariate Euclidean distance of each cell to its nearest untreated Jurkat T-cell neighbors according to their cell cycle molecular profile (**Figure 5E**). NN analysis captures noncanonical cells with skewed phenotypic distance in treatment conditions compared to WT cells (**Figure 5F**). Using regression analysis, we model the euclidean aberrancy score to identify aberrantly expressed features from each treatment. Discovered features that represent noncanonical CC states include increased DNA content reflecting mitotic slippage of G0G1 phase NOC-treated cells; decreased pRb (S780) in G2 phase PALBO- and NOC-treated cells but M phase PALBO- and HU-treated cells; Ki67 loss in S phase PALBO- and NOC-treated cells indicating proliferative aberrancy; and increased PLK1 expression in G0G1 phase NOC-treated cells, and a loss of PLK1 in G2 phase PALBO-treated cells (**Figure 5G**). These noncanonical skewed cells are clearly observed in lower dimensional CC embeddings of each CC phase (**Figure S6F**). Considering these are cell lines with demonstrated noncanonical CC states, canonical cells defined using the NN distance threshold that still demonstrate skewed phenotypes from the untreated setting exist because of short Euclidean distances to natural-occurring noncanonical cell states in the untreated cells.

Finally, we integrate protein biosynthesis analysis using puromycin incorporation and detection with our scCC platform to further understand whether noncanonical cells induced by drug treatment affects protein biosynthesis. Typically, in line with what we have previously reported, we found *de novo* protein synthesis active in G0G1 through G2/M (8). Here, we observe protein translation is reduced because of CC inhibitor treatments, particularly in NOC-treated cells (**Figure S6G**) with affected cells enriched in all stages except S-phase. This loss in protein translation is skewed among noncanonical CC states induced by treatment that are phenotypically more distant from WT cells (**Figure S6H**). Taken together, our scCC approach not only captures the landscape of CC states that arise from pharmacologic inhibition but also flexibly integrates with other scMC technologies like *de novo* protein synthesis detection to reveal the link between protein translation and CC inhibitor action.

### Partitioning variance from cell lines, cell cycle phase, treatments, and CC aberrancy using mixed modeling

Considering our platform detects single-cell heterogeneity across cell line, cell cycle phase, CC inhibitors, and CC aberrancy, we generated a massively parallel single-cell dataset of multiple cell line replicates and drugs in each cell line to quantify the main and interaction effects across cell line identity, drug treatments, CC phase, and CC aberrancy using mixed effects modeling **(Figure 5H, S7A-C)**. Variance partitioning revealed that cell line identity drives the most differences, followed by cell cycle phase, and moderate contribution due to cell cycle aberrancy or treatment effects that interact with cell line identity **(Figure 5I)**. For all cell cycle measurements, we binned variance explained in each coefficient into <5%, 5-10%, 10-20%, and >20%. Notably many transcription factors involved in CC regulation like RUNX1, EZH2, TCF1, FoxM1, and Jarid2 were partially explained by CC aberrancy **(Figure 5J, S7B)**. Cell-line dependent drug sensitivity was detected in DNM1, FoxM1, WGA, EZH2, pRb, SLBP, pCDC2, and DNA content. Molecules like SLBP, PCNA, p-CDC2, pRb, MLL1, and *de novo* protein synthesis also demonstrated cell-line dependent variance with treatment or phase. PLK1, DNMT1, and EZH2 had partially explained variance by interactions across cell line, phase, and aberrancy. This analysis demonstrates the complexity of how these different biological factors are layered to create heterogeneity in cell cycle states. IdU and pH3(s10) were used to manually discretize S and M phase cells for analysis and had high variance for cell cycle gates, which may mask their contribution to other variables. Notably many transcription factors involved in CC regulation like RUNX1, EZH2, TCF1, FoxM1, and Jarid2 were partially explained by CC aberrancy **(Figure 5J, S7D-E)**. Molecules like SLBP, PCNA, phospho-CDC2, and *de novo* protein synthesis also demonstrated cell-line dependent variance with treatment and phase **(Figure S7B)**. Variance partition results as well as differential abundance analysis are shared in **Tables S6-8**. In summary, our platform enabled us to quantify the effects of different biological factors involved in scCC state diversity.

### Assessing Inhibitor and CC States on Ex Vivo Stimulated Primary Human T Cells

Although the induction of aberrant cells with noncanonical CC states using CC inhibitors is an observed phenomenon with adherent cells using microscopy, transformed cell systems like tumor cell lines have documented aberrant CC behaviors compared to healthy primary cells. Thus, the induction of aberrant cells with noncanonical CC states using CC inhibitors may be exacerbated by CC perturbation (46). Healthy primary human T cells from the peripheral blood are majority quiescent suspension cells that undergo rapid expansion upon activation and have been a recent focus of ex vivo expansion efforts for their various clinical applications, emphasizing the importance of deeper CC analysis to understand the therapeutic relevance of expanded cell products (52). Thus, deeper CC phenotyping may help us understand how different drugs affect the same system as well as the canonical and noncanonical CC behaviors that arise in healthy T cell settings. To better characterize the effects of CC inhibition on normal and aberrant CC states in healthy cell systems, we use *ex vivo* TCR stimulation of primary human T cells treated with various CC inhibitors from Day 0-3 of activation (**Figure 6A**). All drugs achieve CC arrest after three days of stimulation that more closely resembles the CC state of D0 unstimulated T cells than the D3 cells, except for Ro-3306, which still demonstrates inhibitor action by a reduction in the abundance of DNA licensing factor CDT1 (**Figure 6B-C**).

**Figure 6.**
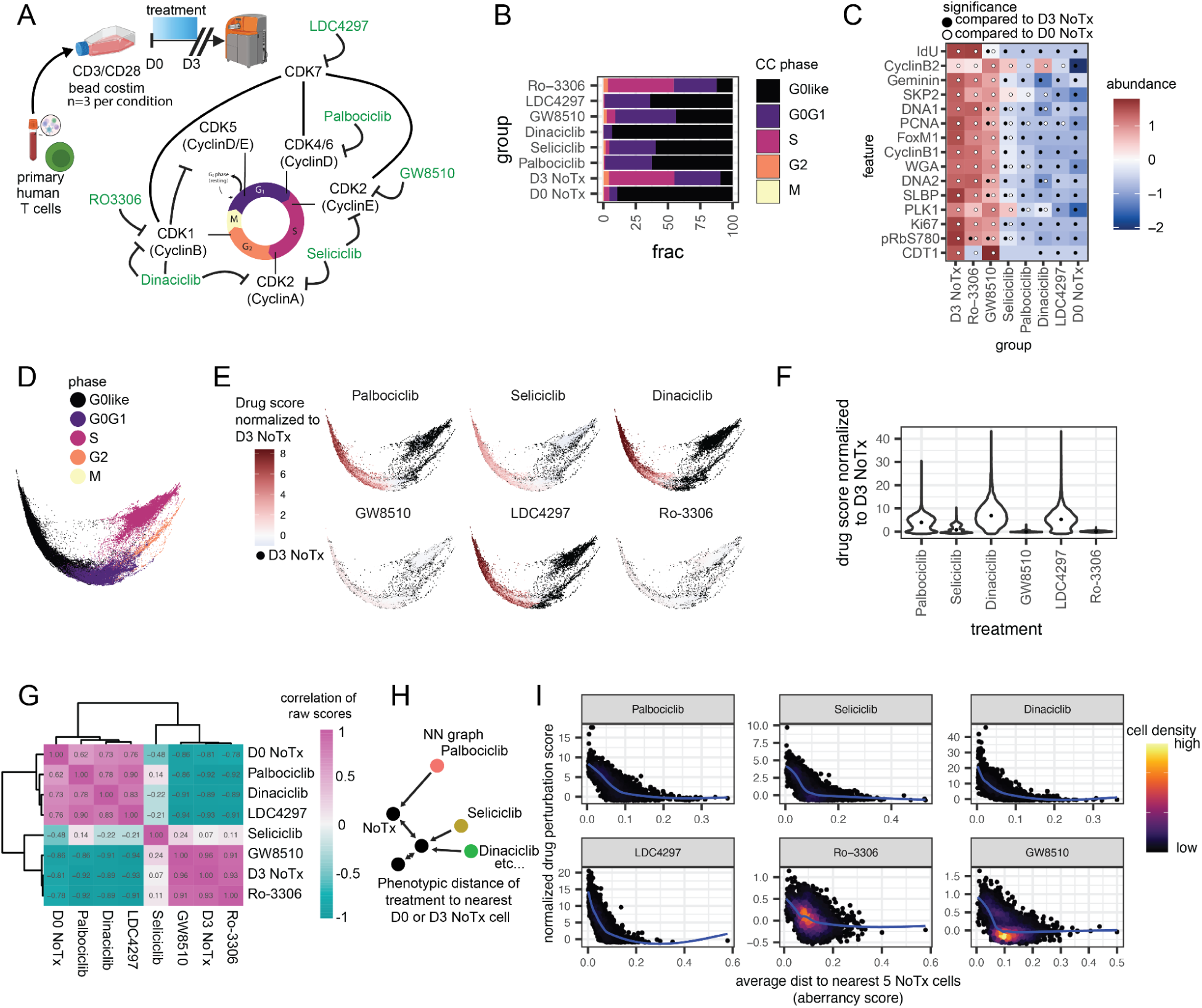
CC aberrancies are associated with escape from CC inhibitor action on ex vivo stimulated primary human T cells. **(A)** Cartoon schematic of experimental design using ex vivo stimulated primary human T cells from healthy donor with various selective CDK inhibitors (n=3 per condition). **(B)** Example CC phase fraction for each treatment. **(C)** Median abundance of CC features. Significance defined by padj<=0.1 and coefficient difference > 0.5 standard deviations. Black dots indicate significance compared to D3 NoTx, and white dots indicate significance compared to D0 NoTx. **(D)** CC embedding of all cells colored by group and **(E)** colored by drug score normalized to NoTx (MELD) for each treatment. **(F)** Relative drug scores for each treatment normalized to D3 NoTx. **(G)** Correlation between raw drug treatment scores (padj<=0.05 for all correlations except Seliciclib). **(H)** Cartoon schematic of NN analysis. **(I)** Drug perturbation score and CC aberrancy score for each treatment colored by cell density.

### Primary human T cells that escape CC inhibitor action have more dissimilar CC phenotypes to untreated cells

To further understand the effects of these drugs on scCC states in healthy human settings, we quantify CC state aberrancy and inhibitor action using NN analysis. Similar to cell lines, we observe an enrichment of noncanonical CC states as a consequence of pharmacologic inhibition of CC (**Figure S8A**). For example, Ki67 loss in G2 cells, a reduction in DNA replication measured using IdU incorporation in LDC4297 and Palbociclib, a reduction in PCNA for Dinaciclib and Palbociclib, and PLK1 reduction in M cells across all treatments. Drug perturbation scores normalized to the untreated D3 cells shows Palbociclib, Seliciclib, Dinaciclib, and LDC4297 with strong arrest signatures at G0like and G0G1 CC states (**Figure 6C-F, Figure S8B**). Correlative analysis of raw scores to compare single-cell inhibitor action between the different treatments shows that Palbociclib, Dinaciclib, LDC4297, and Day0 NoTx CC states are strongly correlated with each other, and Seliciclib correlations are distinct from other treatments (**Figure 6G**). Despite the strong correlations between Palbociclib, Dinaciclib, and LDC4297, these correlations are not perfect and there remains some CC state diversity that is unique to each drug. While GW8510 and Ro-3306 did have a minor effect on CC states, the drug CC states were still strongly correlated with D3 NoTx cells. Notably, normalizing inhibitor action for each cell to the D3 NoTx score shows that the cells affected with shared signatures with each drug were strongly correlated, with some unique CC state diversity as a result of each inhibitor (**Figure S8C-F**). To better understand this CC state diversity, we further analyze the aberrancy of cells that escape inhibitor action by quantifying phenotypic distance of CC states for all cells to any untreated cell at Day 0 or Day 3 (**Figure 6H**). Aberrancy analysis revealed that cells escaping inhibitor action have CC states that are disparate from untreated cells, and is more evident in cells that have S and G2 CC signatures (**Figure 6I, Figure S8G**). Taken together, this analysis suggests that while CDK inhibitors do induce some unique diversity in CC states during *ex vivo* TCR stimulation of healthy primary human T cells, they have globally similar inhibitor action, and that cells escaping arrest demonstrate more aberrant CC states.

In summary, our scCC approach reveals the utility of combining deep scCC phenotyping with pharmacologic inhibition to guide our understanding of how different drugs affect a system. By looking specifically within quiescent healthy human primary cells that proliferate only upon stimulation, we can study the canonical and noncanonical behaviors induced by CC inhibition in a system where cell proliferation and expansion is relevant for producing therapeutic cell products. Our results also suggest that pharmacologic control of CC may be used to tune proliferation during the stimulation phase of manufacturing therapeutic cell products.

## Discussion

The effects of CC state and proliferation pervades diverse systems like cell differentiation, cellular function, and cancer. As technological advances in single-cell proteomic measurement capabilities expand and enable multi-systems analysis, there is a need to increase measurements of CC biology to more deeply characterize CC directly along with the crosstalk between CC and other systems. Here we greatly extended existing approaches for cytometric CC analysis by capturing the expression of new proliferation molecular regulators, licensing factors, CC checkpoint inhibitors, and chromatin states. By combining this molecular panel with barcoding strategies to increase throughput and control for technical effects, we demonstrate the large-scale capture and quantification of diverse cell cycle states across millions of cells and multiple cell lines with pharmacological perturbation. We further demonstrated the generalizability of our molecular panel in CC-arrested primary human T cells.

While CC progression is controlled by the synthesis and degradation of cyclins in an oscillatory manner, CC checkpoints serve to regulate this molecular orchestra during error. Our understanding of canonical CC progression has been historically shaped by bulk assays like western blots or low throughput, low-dimensional imaging assays (12,53). These canonical definitions inform normal CC progression but do not fully capture the single-cell landscape of CC states. With the advent of sc technologies, CC remains coarsely studied as the major phases, CC signatures are often regressed out instead of directly studied, is limited by adherent systems from imaging platforms, or CC biology is inadequately captured because it is fundamentally a protein- and PTM-mediated process, leaving much to be desired for deep CC phenotyping in sc assays like scRNA-seq (54,55). However, aberrant CC states are a reported phenomenon in perturbation systems like microtubule destabilization and cancer is known to induce noncanonical CC states (50,56). Using our CC molecular panel, we capture the diverse single-cell CC states naturally abundant in cell lines that do not qualify canonical, discretized CC states defined in the literature as well as pharmacologically-induced noncanonical CC states. For example, Ki67–often used to identify proliferative cells–was a dynamic molecule with low expression in a subset of Jurkat cells with other molecular signatures of proliferation but also decreased across all phases in the presence of CDK4/6 inhibition. CyclinB2, but not CyclinB1, increased in G0G1 cells with all treatments, which is particularly interesting because both activate CDK1 though CyclinB2 is thought to have a less important role for viability but can compensate for CyclinB1 (57). SLBP, typically expressed in S phase, was increased in G0G1 and G2 cells treated with HU. Importantly, we’ve demonstrated that our scCC platform captures these nuanced cell abundances, is amenable to largescale high-throughput suspension and adherent cells, and can be easily integrated with other cell profiling approaches, all of which can be easily extended to dissociated tissues.

While we observed noncanonical cells naturally existing in cell culture systems, cell lines are transformed with atypical proliferative behaviors, if not already derived from cancer cells (58–60). Thus, it is possible that the induction of noncanonical CC states is an artifact of transformed systems known to have CC aberrancies. To answer this, we further assessed the effectiveness of diverse CDK inhibitors that target a range of CDK molecules controlling different cell cycle checkpoints in CD3/CD28 costimulated primary human T cells from healthy donors. While CDK inhibitors had quantifiable differences in the induced scCC states, the effects of CDK inhibition primarily resulted in G0G1 arrest and drug action was strongly correlated for Palbociclib, Dinaciclib, and LDC4297. Notably, primary human T cells that escaped inhibitor action had more phenotypically aberrant CC states when compared to untreated T cells, demonstrating that noncanonical CC states are pharmacologically inducible in healthy systems as well. Our understanding of noncanonical CC states will continue to evolve as we more deeply understand the dynamics and roles of CC molecules in healthy and perturbation settings.

In summary, our CC-related molecular panel can capture the diversity of canonical and noncanonical scCC biology in both cell line and primary cell systems, which can be further integrated with perturbation systems to dissect granular CC biology. This is consistent with previous single-cell transcriptomic studies showing that CC drug-tolerant PC9 cells occupy heterogenous and distinct cell subpopulations and cell states (61). These distinct cells that escape early CC arrest and survive may also be noncanonical seed cells for drug tolerant cells. In the case of primary cells, cell proliferation is an essential metric for therapeutic cell products. Previous studies demonstrate that cell proliferation and cell fate are tightly linked in T cells, where tuning TCR stimulation has a direct impact on CC progression and division (26). Extending our scCC platform to T cells, we demonstrated that CC slowing with CDK inhibition has direct consequences on CC state, which future studies can harness to understand whether CC tuning is useful for the manufacturing process of cell therapeutics.

We present a robust and generalizable platform for parallel scCC measurements using CyTOF to deeply characterize CC states that future studies can leverage in tandem with other mass cytometry molecular panels for sc and dissociated tissue systems. Multiplexing our scCC panel with published or custom panels such as epigenetic state using EpiTOF, chromatin content using Chromotyping, single-cell metabolic regulome profiling (scMEP), immune cell monitoring, or cell differentiation in perturbation and disease settings can be valuable to dissect the crosstalk between CC and other cellular processes in different areas of life science research.

## Methods

### Data availability

All data (fcs files) are ready for public release. Any additional information desired in this article is available from the lead contact upon request.

### Cell line culture

Five cell lines were used in this study: NALM6, U937, HEL, 293T (ATCC CRL-3216), JURKAT (ATCC CRL-2899). NALM6 is a lymphocyte-like cell line derived from human B cell precursor leukemia, U937 is a pro-monocytic cell line originating from human myeloid leukemia, HEL is an erythroblast cell line derived from human erythroleukemia, 293T is an adherent epithelial-like cell line originating from human embryonic kidney cells, and JURKAT is a CD4+ lymphocyte-like cell line derived from human T cell acute leukemia. The NALM6, JURKAT, U937, and HEL cells were maintained in RPMI 1640 (Gibco, 11-879-020), 10% FBS (Sigma-Aldrich, F4135), 1% p/s (Thermo Fisher Scientific, 15140122), and Glutamax (Thermo Fisher Scientific, 35-050-061). 293T cells were maintained in DMEM (Dulbecco’s Modified Eagle’s Medium)/Hams F-12 50/50 Mix (Corning, 10-090-CV), 10% FBS, and 1% p/s. Cells were maintained at 37 °C, 5% CO2.

### Ex vivo Primary human T cell TCR stimulation

De-identified peripheral blood and LRS chambers samples from healthy human donors were obtained, and experiments were carried out following guidelines of the Stanford Institutional Review Board (IRB). Collections were monitored and reviewed by Stanford’s IRB. Written informed consent was obtained from all participants managed by Stanford Blood Center. PBMCs were isolated via Ficoll (GE Healthcare) density gradient centrifugation. Bulk T cells were negatively isolated from whole blood using RosetteSep Human T Cell Enrichment Cocktail (StemCell Technologies). T cells were cultured in ImmunoCult-XF T Cell Expansion Medium (10981, Stemcell Technologies) and supplemented with 10 ng ml−1 of interleukin-2 (Miltenyi Biotec). T cells were activated and expanded using Human T-Expander CD3/CD28 (Dynabeads, Thermo Fisher) added in a 1:1 cell-to-bead ratio and cells were incubated at 37 °C in 5% CO2.

### IdU labeling

5-Iodo-2-deoxyuridine (Sigma I7125) was re-suspended in DMSO (Sigma D2650) at 500 mM. IdU labeling was performed at a final concentration 100 μM and returned to the incubator for 15-30 minutes. After labeling, cells were washed with PBS and continued with the live-dead labeling and fixation protocol for CyTOF.

### Live/dead labeling, cell fixation, and permeabilization for CyTOF staining

CyTOF staining was performed as previously described, following this protocol.io publication (62). Briefly, live-dead labeling was performed using cisplatin in low-barium PBS, cells were washed with CSM, fixed with PFA for 10 minutes at room temperature, and stored in -80C until staining. When ready, fixed samples were thawed at room temperature and barcoded using fixed-cell palladium barcoding, pooled into a single tube. Surface staining was performed with metal-conjugated antibodies in CSM for 30 minutes at room temperature, washed, permeabilized with 100% 4c methanol and incubated on ice for 10 minutes, washed 3x times with CSM, and proceeded with intracellular staining. Finally, cells were washed with CSM, and resuspended in an intercalator solution until CyTOF analysis.

### Drug treatments

Drug treatments were performed using ibrutinib (PCI-32765; Cellagen Technology #C7327), palbociclib (Chemscene, CS-3110), roscovitine (Seliciclib, CYC202, Selleck Chemicals #S1153), dinaciclib (SCH727965, Selleck Chemicals, #S2768), LDC4297 (LDC044297, Selleck Chemicals, #S7992), ro3306 (Selleck Chemicals,#7747), GW6510 (sc-biotech #sc-215122), hydroxyurea (Sigma-Aldrich #H8627-1G), nocodazole (Sigma-Aldrich. #SML1665-1ML), SU9516 (EMD Millipore, Calbiochem, #572650). Concentrations are provided in Table S5.

### Palladium barcoding and staining with metal-conjugated antibodies

Individual samples within one experiment were palladium barcoded as described previously (14) and combined into a single sample before further processing and staining. Experiments with multiple barcode plates had an anchor sample to normalize any technical effects.

### CyTOF processing

Raw mass cytometry data were bead normalized to remove acquisition-related influences on marker expression using the premessa R package. Sample barcoding was done using fixed cell palladium barcode combinations and debarcoded using premessa. Normalized data were uploaded to CellEngine for bead removal, singlet identification, removal of debris and non-biological events, live cell gating, and exported (https://www.cellengine.com/). Pre-apoptotic cells defined by cPARP positivity were also removed. Batch correction between multiple palladium barcodes in the same experiment was performed using an anchor sample with an adjustment factor for each channel (63). Adjusted data was imported into CellEngine for manual cell cycle gating using CyclinB1, IdU labeling, and phospho-HistoneH3 (s10). Gated files were subsequently imported into the R environment, asinh transformed (cofactor 5), and normalized to the 99.9th percentile of each respective channel before downstream analysis.

### Dimensionality reduction, quantifying perturbations, and diversity

We used PHATE (R/Python) with knn_dist=cosine, mds_dist=euclidean, and knn=15. We used RANN::nn2 in R or sklearn’s KNeighborsClassifier in Python to find the nearest WT or untreated cell neighbors by euclidean or cosine distance. Single-cell perturbation scores were quantified using MELD with default parameters. Analysis and plotting was done in R v4.2. For cell state diversity, we downsampled to 2,000 cells per cell line and used Ki67, pRbS780, pH3_s10, CDT1, IdU, Geminin, PLK1, DNA, CyclinB1, PCNA, SLBP, CyclinE, WGA, HH3, EZH2, CTCF, and MLL1. We computed an adjacency matrix for all possible combinations of features using a cosine distance with a threshold of 0.5 and quantified graph density using igraph. Statistical models were computed using a random, balanced sampling of 50,000 cells per cell line.

### Statistical analysis

We use the GLM/M framework in diffCyt for statistical comparisons unless otherwise noted in the figure legends. Multiple hypothesis correction was performed in R using p.adjust() with a False Discovery Rate (benjamini hochberg) correction procedure. Proportion of variance calculations for cell line associations were performed using lm and anova in R, the sum of squares was calculated, and the proportion of variance explained was computed from the total sum of squares. Multivariate logit models were computed using nnet::multinom in R, and pairwise sensitivity, specificity, and accuracy were calculated using pROC::multiclass.roc. Variance partitioning was performed using the variancePartition R package and including main effects as well as all two-way, three-way, and four-way interaction effects across cell lines, drug treatment, aberrancy, and cell cycle phase.

## Acknowledgements

We thank Jolene Ranek for manuscript comments and suggestions, and Simone de Beauvoir for her canine support in writing this manuscript.

## Contributions

Conceptualization, MA and SCB. Methodology, MA. Software, MA. Formal analysis, MA. Resources, MF, PF, TB, DH, SCB. Data curation, MA. Writing, MA and SCB. Editing, MA, SCB, PF.

## Supplementary Figures: Amouzgar et al. 2025

**Figure S1:**
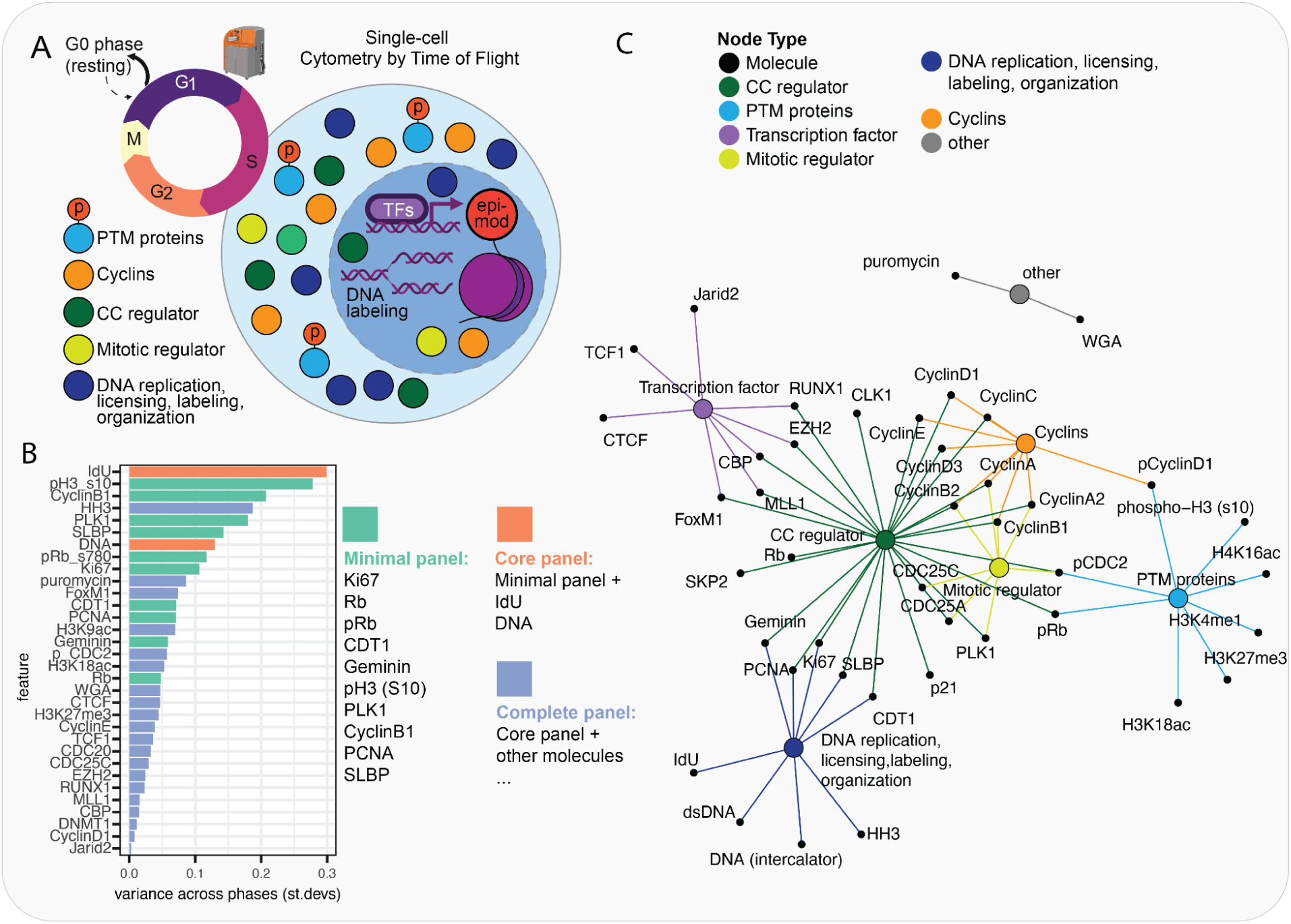
Cartoon diagram of CC related molecules mapped to different categories. **(A)** Measured CC molecules are mapped to different categories. CC regulator=direct control of CC progression. PTM proteins=post translational modification. Mitotic regulator=direct control of G2M transition or mitotic progression. DNA replication, licensing, labeling, organization=measurements relating to DNA replication during S phase, DNA replication licensing, nucleotide labeling, intercalation, or histone content. Cyclins=Cyclin molecule, phosphorylated or not. Transcription factor=Any molecule also categorized as a transcription factor. Other=WGA for cell size, and puromycin for de novo protein synthesis measured using puromycin incorporation. **(B)** Feature variance across cell cycle phases to detect most variable targets for separating cell cycle progression. **(C)** Graphical diagram of features and category mapping.

**Figure S2:**
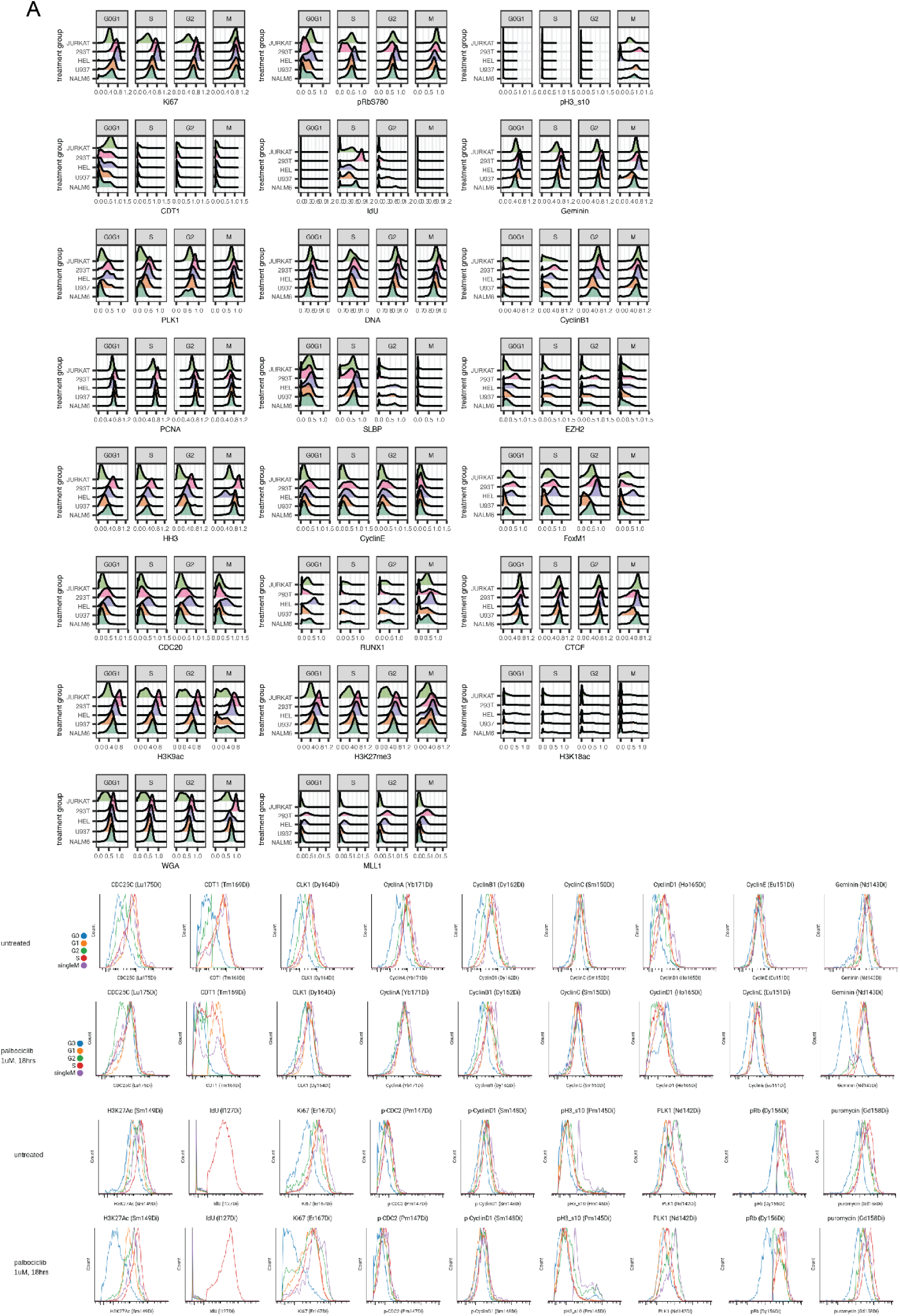
Molecular abundance of CC related targets across cell lines. **(A)** Smooth histograms of example CC targets across cell lines and drug treatments.

**Figure S3:**
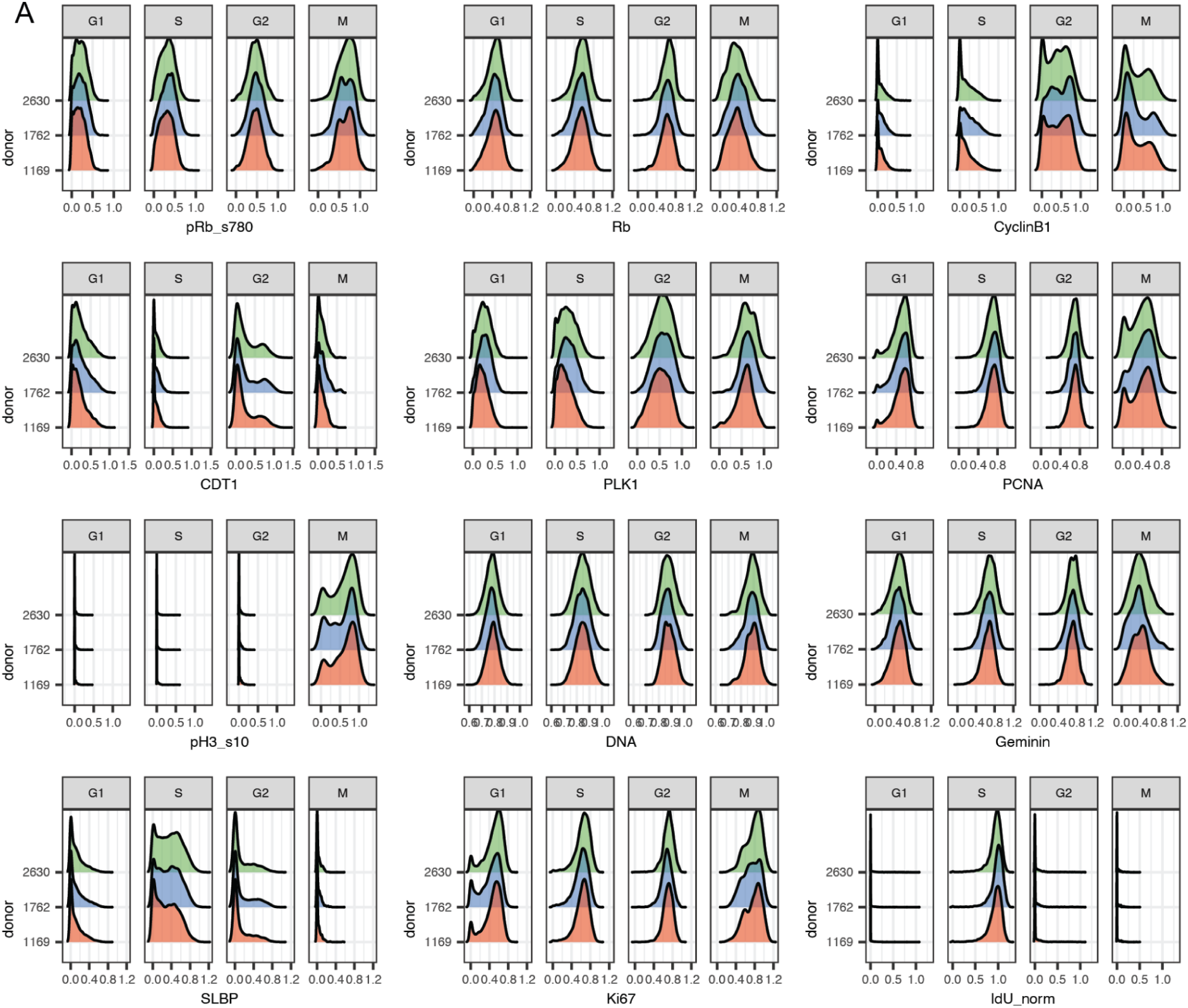
Molecular abundance of core CC targets across 3 primary human T cells from 3 donors. (A) Smooth histograms of CC markers across each donor. **(A)** CC fractions across different cell line replicates. **(B)** Hierarchically clustered heatmap of mean cell cycle molecular abundance across cell line replicates. **(C)** Differential abundance analysis of all pairwise cell lines for different cell cycle related markers with # of significant features (barplot), scaled model coefficient estimate aka change in abundance (heatmap), and significance labels defined by padj<=0.1 and estimates >= 0.25 (points). **(D)** CC embedding before and after integrating cell lines using harmony. **(D)** CC phase purity analysis of clustering before and after correction. **(E)** FlowSOM with 4 clusters colored by cell cycle phase (top) and **(F)** Fraction of each cell line in each cluster (equally sampled across cell lines, bottom). **(G-I)** Variant analysis comparing CC genes for cell lines across three scales of variant information available for Jurkat, HEL, U-937, and NALM-6 but not HEK293T in the DepMap database: (1) exact SNV (most specific), (2) variant type in a gene regardless of specific nucleotide position (eg, a missense variant in ABR regardless of specific SNV), and (3) CNVs. **(G)** Upset plot of exact SNVs across different cell lines. 1 unique missense SNV shared between Jurkat and NALM-6 cells: a Guanine to Adenine nucleotide change in ABR at ENST00000302538.10:c.2285G>A. **(H)** Binary heatmap of variant type in a gene regardless of specific nucleotide position. **(I)** Heatmap of genes binned for increased, no change, or decreased CNVs detected in a cell line.

**Figure S4:**
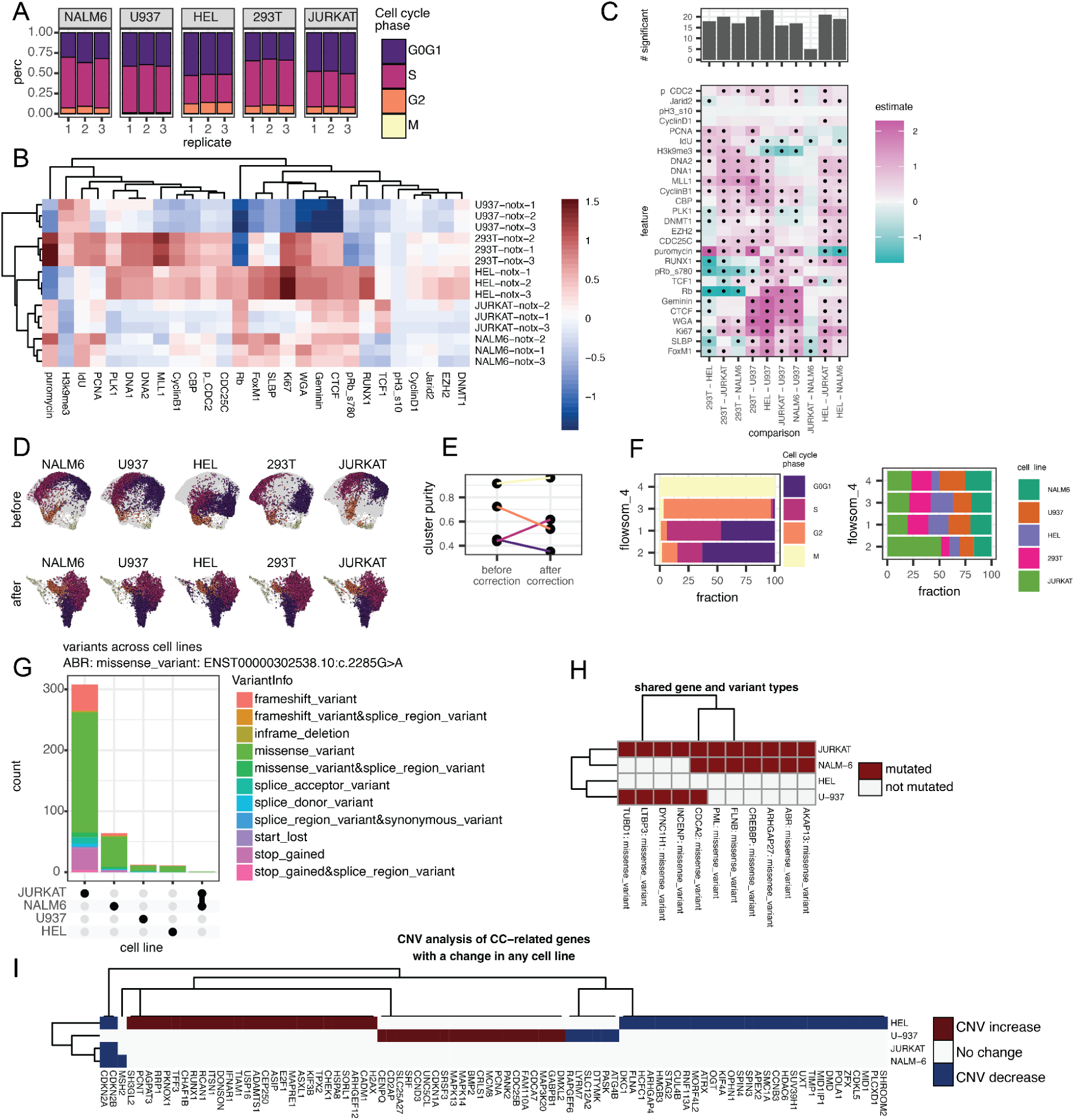
Canonical and noncanonical CC states across cell lines. **(A)** CC fractions across different cell line replicates. **(B)** Hierarchically clustered heatmap of mean cell cycle molecular abundance across cell line replicates. **(C)** Differential abundance analysis of all pairwise cell lines for different cell cycle related markers with # of significant features (barplot), scaled model coefficient estimate aka change in abundance (heatmap), and significance labels defined by padj<=0.1 and estimates >= 0.25 (points). **(D)** CC embedding before and after integrating cell lines using harmony. **(D)** CC phase purity analysis of clustering before and after correction. **(E)** FlowSOM with 4 clusters colored by cell cycle phase (top) and **(F)** Fraction of each cell line in each cluster (equally sampled across cell lines, bottom). **(G-I)** Variant analysis comparing CC genes for cell lines across three scales of variant information available for Jurkat, HEL, U-937, and NALM-6 but not HEK293T in the DepMap database: (1) exact SNV (most specific), (2) variant type in a gene regardless of specific nucleotide position (eg, a missense variant in ABR regardless of specific SNV), and (3) CNVs. **(G)** Upset plot of exact SNVs across different cell lines. 1 unique missense SNV shared between Jurkat and NALM-6 cells: a Guanine to Adenine nucleotide change in ABR at ENST00000302538.10:c.2285G>A. **(H)** Binary heatmap of variant type in a gene regardless of specific nucleotide position. **(I)** Heatmap of genes binned for increased, no change, or decreased CNVs detected in a cell line.

**Figure S5:**
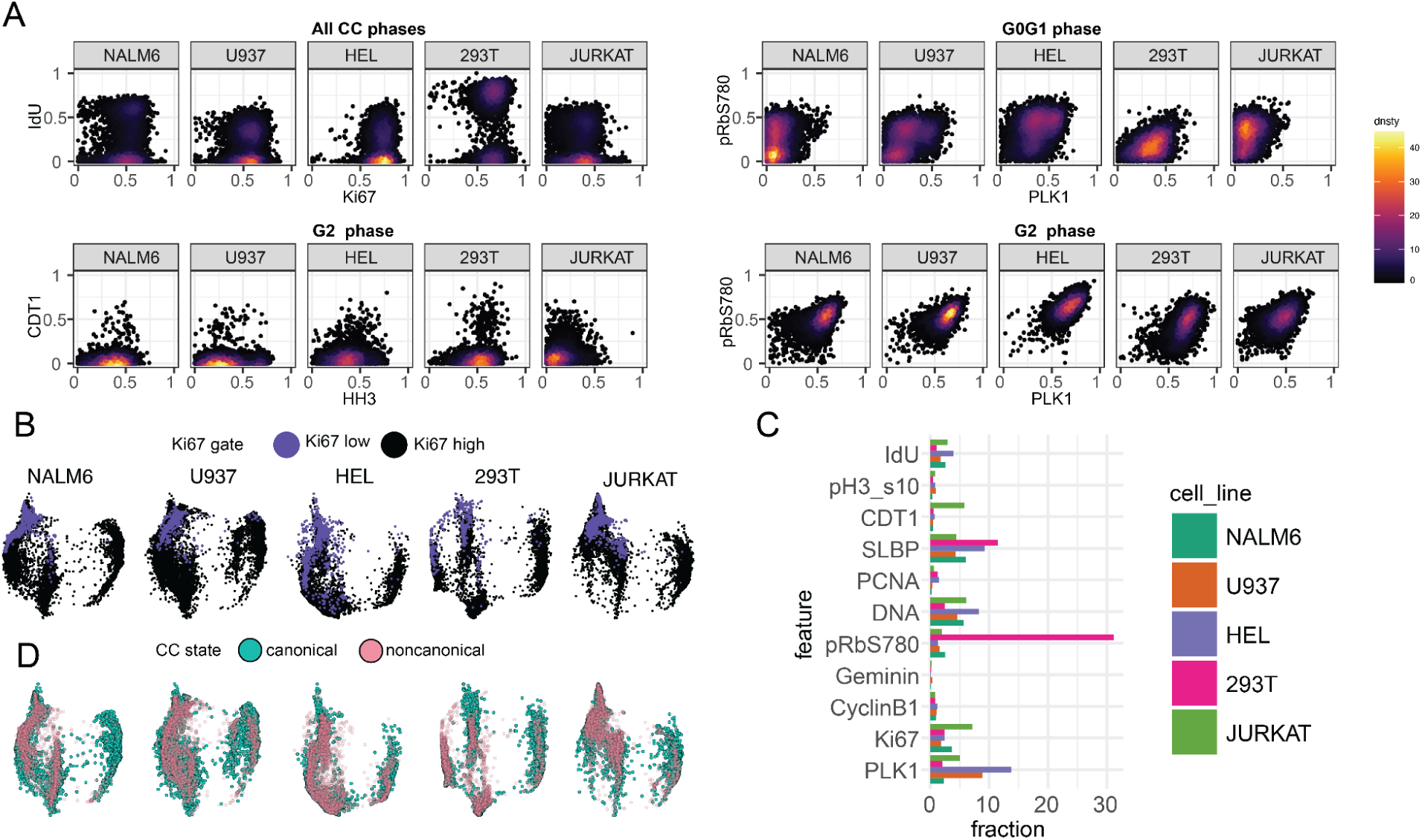
Canonical and noncanonical CC states across cell lines. **(A)** Flow plots with example markers for phases with noncanonical states. **(B)** CC embeddings for each cell line colored by Ki67 low abundance. **(C)** fraction of cells with noncanonical identities based on manual discretization in each cell line. **(D)** CC embeddings for each cell line colored by canonical or noncanonical state.

**Figure S6:**
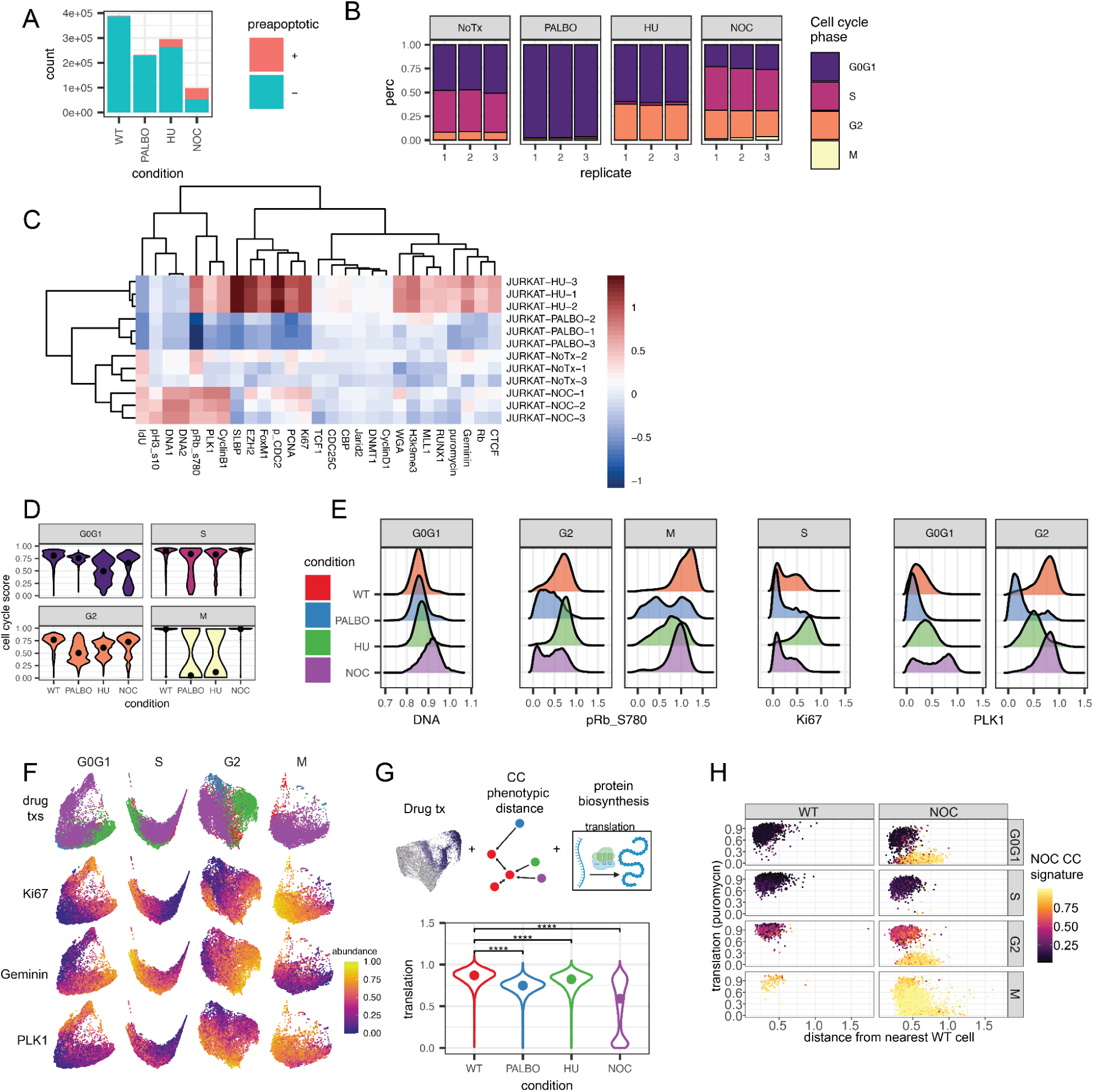
Drug perturbation effects. **(A)** Removal of cPARP+ pre-apoptotic cells. # of cells before gating; WT: 390074, PALBO: 232619, NOC: 97727, HU: 295362. Remaining normal cells after removal: WT: 386,643, PALBO: 229,629, HU: 263,428, NOC: 53,129. **(B)** Cell cycle fractions for 3 replicates of each drug in Jurkat cells. CC embeddings faceted by treatment condition. **(C)** Heatmap of molecular abundance for all replicates. **(D)** Cell cycle phase scores. **(E)** Smoothed histograms for example markers of noncanonical cells. **(F)** CC embeddings of canonical and noncanonical cells computed on each group with example markers. **(G)** Cartoon schematic integrating drug perturbations with phenotypic distance analysis for noncanonical CC states and protein biosynthesis measurements using MC and drug perturbation results on protein biosynthesis. **(H)** Relationship between protein translation, phenotypic distance, and NOC signature for NOC-treated and WT cells.

**Figure S7:**
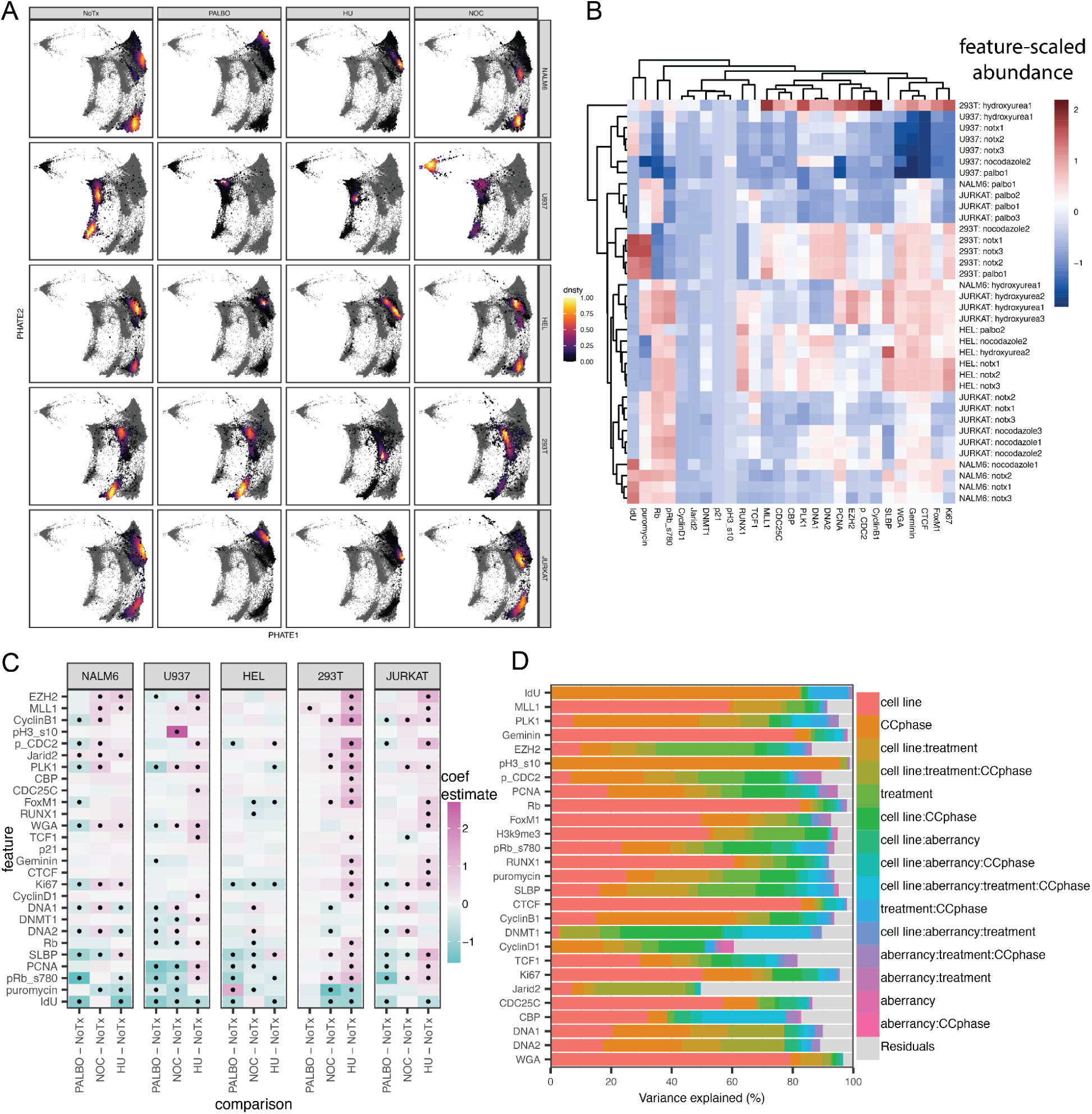
Drug perturbation effects. **(A)** CC embedding of cell lines from NoTx, Palbociclib, Hydroxyurea, and Nocodazole treatments. **(B)** Heatmap of all replicates for cell lines from NoTx, Palbociclib, Hydroxyurea, and Nocodazole treatments. **(C)** Heatmap of coefficient estimates (fold-change) comparing drug treatment to no treatment in each cell line. Significant features are detected with an adjusted-pvalue threshold <=0.1, and minimum absolute value coefficient (fold-change) threshold of 0.25 **(D)** Percentage of variance explained for all main and interaction effects in each feature.

**Figure S8.**
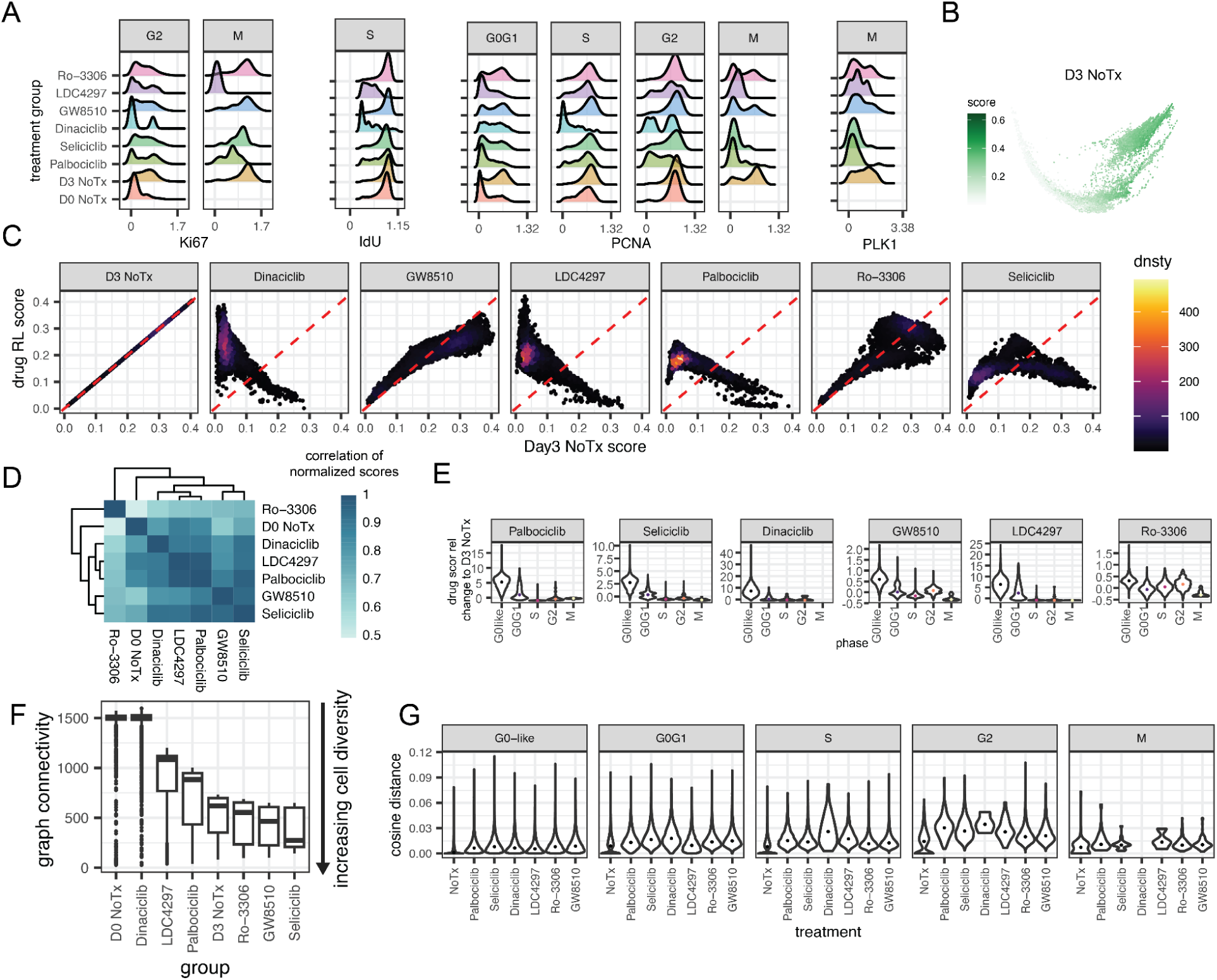
CC inhibitor action on ex vivo stimulated primary human T cells. **(A)** Expression of example CC targets for each condition in different groups. **(B)** CC embedding colored by D3 NoTx score. **(C)** D3 NoTx likelihood score versus each drug’s likelihood score. **(D)** Correlation between drug treatment effects. **(E)** Normalized drug likelihood score across each phase. **(F)** Cell diversity analysis for each treatment using graph connectivity analysis. **(G)** CC phenotypic distance for each treatment across each phase.

## References

1. Wiman KG, Zhivotovsky B. Understanding cell cycle and cell death regulation provides novel weapons against human diseases. J Intern Med. 2017 May;281(5):483–95.

2. Laphanuwat P, Jirawatnotai S. Immunomodulatory Roles of Cell Cycle Regulators. Front Cell Dev Biol. 2019;7:23.

3. Chong KH, Zhang X, Zheng J. Dynamical analysis of cellular ageing by modeling of gene regulatory network based attractor landscape. PloS One. 2018;13(6):e0197838.

4. Vignon C, Debeissat C, Georget MT, Bouscary D, Gyan E, Rosset P, et al. Flow cytometric quantification of all phases of the cell cycle and apoptosis in a two-color fluorescence plot. PloS One. 2013;8(7):e68425.

5. McKinnon KM. Flow Cytometry: An Overview. Curr Protoc Immunol. 2018 Feb 21;120:5.1.1–5.1.11.

6. Bendall SC, Simonds EF, Qiu P, Amir E ad D, Krutzik PO, Finck R, et al. Single-cell mass cytometry of differential immune and drug responses across a human hematopoietic continuum. Science. 2011 May 6;332(6030):687–96.

7. Bandura DR, Baranov VI, Ornatsky OI, Antonov A, Kinach R, Lou X, et al. Mass cytometry: technique for real time single cell multitarget immunoassay based on inductively coupled plasma time-of-flight mass spectrometry. Anal Chem. 2009 Aug 15;81(16):6813–22.

8. Kimmey SC, Borges L, Baskar R, Bendall SC. Parallel analysis of tri-molecular biosynthesis with cell identity and function in single cells. Nat Commun. 2019 Mar 12;10(1):1185.

9. Behbehani GK. Cell Cycle Analysis by Mass Cytometry. Methods Mol Biol Clifton NJ. 2018;1686:105–24.

10. Rein ID, Notø HØ, Bostad M, Huse K, Stokke T. Cell Cycle Analysis and Relevance for Single-Cell Gating in Mass Cytometry. Cytometry A. 2020 Aug;97(8):832–44.

11. Pack LR, Daigh LH, Meyer T. Putting the brakes on the cell cycle: mechanisms of cellular growth arrest. Curr Opin Cell Biol. 2019 Oct;60:106–13.

12. Stallaert W, Taylor SR, Kedziora KM, Taylor CD, Sobon HK, Young CL, et al. The molecular architecture of cell cycle arrest. Mol Syst Biol. 2022 Sep;18(9):e11087.

13. Hartmann FJ, Simonds EF, Bendall SC. A Universal Live Cell Barcoding-Platform for Multiplexed Human Single Cell Analysis. Sci Rep. 2018 Jul 17;8(1):10770.

14. Zunder ER, Finck R, Behbehani GK, Amir EAD, Krishnaswamy S, Gonzalez VD, et al. Palladium-based mass tag cell barcoding with a doublet-filtering scheme and single-cell deconvolution algorithm. Nat Protoc. 2015 Feb;10(2):316–33.

15. Sakaue-Sawano A, Kurokawa H, Morimura T, Hanyu A, Hama H, Osawa H, et al. Visualizing spatiotemporal dynamics of multicellular cell-cycle progression. Cell. 2008 Feb 8;132(3):487–98.

16. Uzman A. Molecular biology of the cell (4th ed.): Alberts, B., Johnson, A., Lewis, J., Raff, M., Roberts, K., and Walter, P. Biochem Mol Biol Educ. 2003 Jul;31(4):212–4.

17. Satyanarayana A, Kaldis P. Mammalian cell-cycle regulation: several Cdks, numerous cyclins and diverse compensatory mechanisms. Oncogene. 2009 Aug 20;28(33):2925–39.

18. Zatulovskiy E, Zhang S, Berenson DF, Topacio BR, Skotheim JM. Cell growth dilutes the cell cycle inhibitor Rb to trigger cell division. Science. 2020 Jul 24;369(6502):466–71.

19. Zhang H. Regulation of DNA Replication Licensing and Re-Replication by Cdt1. Int J Mol Sci. 2021 May 14;22(10):5195.

20. Driscoll DL, Chakravarty A, Bowman D, Shinde V, Lasky K, Shi J, et al. Plk1 inhibition causes post-mitotic DNA damage and senescence in a range of human tumor cell lines. PloS One. 2014;9(11):e111060.

21. Shi Q, King RW. Chromosome nondisjunction yields tetraploid rather than aneuploid cells in human cell lines. Nature. 2005 Oct;437(7061):1038–42.

22. Cheung P, Vallania F, Warsinske HC, Donato M, Schaffert S, Chang SE, et al. Single-Cell Chromatin Modification Profiling Reveals Increased Epigenetic Variations with Aging. Cell. 2018 May 31;173(6):1385–1397.e14.

23. Hartmann FJ, Mrdjen D, McCaffrey E, Glass DR, Greenwald NF, Bharadwaj A, et al. Single-cell metabolic profiling of human cytotoxic T cells. Nat Biotechnol. 2021 Feb;39(2):186–97.

24. Tsai AG, Glass DR, Juntilla M, Hartmann FJ, Oak JS, Fernandez-Pol S, et al. Multiplexed single-cell morphometry for hematopathology diagnostics. Nat Med. 2020 Mar;26(3):408–17.

25. Hartmann FJ, Babdor J, Gherardini PF, Amir EAD, Jones K, Sahaf B, et al. Comprehensive Immune Monitoring of Clinical Trials to Advance Human Immunotherapy. Cell Rep. 2019 Jul 16;28(3):819–831.e4.

26. Proliferation tracing with single-cell mass cytometry optimizes generation of stem cell memory-like T cells | Nature Biotechnology [Internet]. [cited 2024 Nov 4]. Available from: https://www.nature.com/articles/s41587-019-0033-2

27. Barnum KJ, O’Connell MJ. Cell cycle regulation by checkpoints. Methods Mol Biol Clifton NJ. 2014;1170:29–40.

28. Miller I, Min M, Yang C, Tian C, Gookin S, Carter D, et al. Ki67 is a Graded Rather than a Binary Marker of Proliferation versus Quiescence. Cell Rep. 2018 Jul 31;24(5):1105–1112.e5.

29. Zhu Q, Zhao X, Zhang Y, Li Y, Liu S, Han J, et al. Single cell multi-omics reveal intra-cell-line heterogeneity across human cancer cell lines. Nat Commun. 2023 Dec 9;14(1):8170.

30. Bendall SC, Simonds EF, Qiu P, Amir E ad D, Krutzik PO, Finck R, et al. Single-Cell Mass Cytometry of Differential Immune and Drug Responses Across a Human Hematopoietic Continuum. Science. 2011 May 6;332(6030):687–96.

31. Moon KR, Van Dijk D, Wang Z, Gigante S, Burkhardt DB, Chen WS, et al. Visualizing structure and transitions in high-dimensional biological data. Nat Biotechnol. 2019 Dec;37(12):1482–92.

32. Stern AD, Rahman AH, Birtwistle MR. Cell size assays for mass cytometry. Cytom Part J Int Soc Anal Cytol. 2017 Jan;91(1):14–24.

33. Gillooly JF, Hein A, Damiani R. Nuclear DNA Content Varies with Cell Size across Human Cell Types. Cold Spring Harb Perspect Biol. 2015 Jul 1;7(7):a019091.

34. Rosenbluth MJ, Lam WA, Fletcher DA. Force microscopy of nonadherent cells: a comparison of leukemia cell deformability. Biophys J. 2006 Apr 15;90(8):2994–3003.

35. Röselová P, Obr A, Holoubek A, Grebeňová D, Kuželová K. Adhesion structures in leukemia cells and their regulation by Src family kinases. Cell Adhes Migr. 2018 May 4;12(3):286–98.

36. Shin JH, Lee MG, Choi S, Park JK. Inertia-activated cell sorting of immune-specifically labeled cells in a microfluidic device. RSC Adv. 2014;4(74):39140–4.

37. Korsunsky I, Millard N, Fan J, Slowikowski K, Zhang F, Wei K, et al. Fast, sensitive and accurate integration of single-cell data with Harmony. Nat Methods. 2019 Dec;16(12):1289–96.

38. Luecken MD, Büttner M, Chaichoompu K, Danese A, Interlandi M, Mueller MF, et al. Benchmarking atlas-level data integration in single-cell genomics. Nat Methods. 2022 Jan;19(1):41–50.

39. Zhou W, Koudijs KKM, Böhringer S. Influence of batch effect correction methods on drug induced differential gene expression profiles. BMC Bioinformatics. 2019 Aug 22;20(1):437.

40. Zindler T, Frieling H, Neyazi A, Bleich S, Friedel E. Simulating ComBat: how batch correction can lead to the systematic introduction of false positive results in DNA methylation microarray studies. BMC Bioinformatics. 2020 Jun 30;21(1):271.

41. Amouzgar M, Glass DR, Baskar R, Averbukh I, Kimmey SC, Tsai AG, et al. Supervised dimensionality reduction for exploration of single-cell data by HSS-LDA. Patterns. 2022 Aug;3(8):100536.

42. Zheng L, Dominski Z, Yang XC, Elms P, Raska CS, Borchers CH, et al. Phosphorylation of stem-loop binding protein (SLBP) on two threonines triggers degradation of SLBP, the sole cell cycle-regulated factor required for regulation of histone mRNA processing, at the end of S phase. Mol Cell Biol. 2003 Mar;23(5):1590–601.

43. DepMap B. DepMap 24Q4 Public [Internet]. Figshare+; 2024 [cited 2025 May 19]. p. 30825074613 Bytes. Available from: https://plus.figshare.com/articles/dataset/DepMap_24Q4_Public/27993248/1

44. Hanahan D. Hallmarks of Cancer: New Dimensions. Cancer Discov. 2022 Jan 1;12(1):31–46.

45. Bruinsma W, Aprelia M, García-Santisteban I, Kool J, Xu YJ, Medema RH. Inhibition of Polo-like kinase 1 during the DNA damage response is mediated through loss of Aurora A recruitment by Bora. Oncogene. 2017 Mar;36(13):1840–8.

46. Matson JP, Cook JG. Cell cycle proliferation decisions: the impact of single cell analyses. FEBS J. 2017 Feb;284(3):362–75.

47. Gheghiani L, Loew D, Lombard B, Mansfeld J, Gavet O. PLK1 Activation in Late G2 Sets Up Commitment to Mitosis. Cell Rep. 2017 Jun 6;19(10):2060–73.

48. Wang X, Zhao S, Xin Q, Zhang Y, Wang K, Li M. Recent progress of CDK4/6 inhibitors’ current practice in breast cancer. Cancer Gene Ther. 2024 Sep;31(9):1283–91.

49. Kapor S, Čokić V, Santibanez JF. Mechanisms of Hydroxyurea-Induced Cellular Senescence: An Oxidative Stress Connection? Nayak AK, editor. Oxid Med Cell Longev. 2021 Jan;2021(1):7753857.

50. Blajeski AL, Phan VA, Kottke TJ, Kaufmann SH. G(1) and G(2) cell-cycle arrest following microtubule depolymerization in human breast cancer cells. J Clin Invest. 2002 Jul;110(1):91–9.

51. Burkhardt DB, Stanley JS, Tong A, Perdigoto AL, Gigante SA, Herold KC, et al. Quantifying the effect of experimental perturbations at single-cell resolution. Nat Biotechnol. 2021 May;39(5):619–29.

52. Kalamasz D, Long SA, Taniguchi R, Buckner JH, Berenson RJ, Bonyhadi M. Optimization of human T-cell expansion ex vivo using magnetic beads conjugated with anti-CD3 and Anti-CD28 antibodies. J Immunother Hagerstown Md 1997. 2004;27(5):405–18.

53. Mahdessian D, Cesnik AJ, Gnann C, Danielsson F, Stenström L, Arif M, et al. Spatiotemporal dissection of the cell cycle with single-cell proteogenomics. Nature. 2021 Feb;590(7847):649–54.

54. Guo X, Chen L. From G1 to M: a comparative study of methods for identifying cell cycle phases. Brief Bioinform. 2024 Jan 22;25(2):bbad517.

55. Nestorowa S, Hamey FK, Pijuan Sala B, Diamanti E, Shepherd M, Laurenti E, et al. A single-cell resolution map of mouse hematopoietic stem and progenitor cell differentiation. Blood. 2016 Aug 25;128(8):e20–31.

56. Liu J, Peng Y, Wei W. Cell cycle on the crossroad of tumorigenesis and cancer therapy. Trends Cell Biol. 2022 Jan;32(1):30–44.

57. Bellanger S, De Gramont A, Sobczak-Thépot J. Cyclin B2 suppresses mitotic failure and DNA re-replication in human somatic cells knocked down for both cyclins B1 and B2. Oncogene. 2007 Nov 8;26(51):7175–84.

58. Fukumoto M, Shevrin DH, Roninson IB. Analysis of gene amplification in human tumor cell lines. Proc Natl Acad Sci U S A. 1988 Sep;85(18):6846–50.

59. Little CD, Nau MM, Carney DN, Gazdar AF, Minna JD. Amplification and expression of the c-myc oncogene in human lung cancer cell lines. Nature. 1983 Nov 10;306(5939):194–6.

60. Rangarajan A, Hong SJ, Gifford A, Weinberg RA. Species- and cell type-specific requirements for cellular transformation. Cancer Cell. 2004 Aug;6(2):171–83.

61. Aissa AF, Islam ABMMK, Ariss MM, Go CC, Rader AE, Conrardy RD, et al. Single-cell transcriptional changes associated with drug tolerance and response to combination therapies in cancer. Nat Commun. 2021 Mar 12;12(1):1628.

62. Amouzgar M, Favaro P, Ho D, Bruce T, Vijayaragavan K, Bendall S. CYToF Staining v1 [Internet]. 2024 [cited 2025 Jan 26]. Available from: https://www.protocols.io/view/cytof-staining-dfjb3kin

63. Schuyler RP, Jackson C, Garcia-Perez JE, Baxter RM, Ogolla S, Rochford R, et al. Minimizing Batch Effects in Mass Cytometry Data. Front Immunol. 2019 Oct 15;10:2367.

